# Evolved phenological cueing strategies show variable responses to climate change

**DOI:** 10.1101/436857

**Authors:** Collin B. Edwards, Louie Yang

## Abstract

Several studies have documented a global pattern of phenological advancement that is consistent with ongoing climate change. However, the magnitude of these phenological shifts is highly variable across taxa and locations. This variability of phenological responses has been difficult to explain mechanistically. To examine how the evolution of multi-trait cueing strategies could produce variable responses to climate change, we constructed a model in which organisms evolve strategies that integrate multiple environmental cues to inform anticipatory phenological decisions. We simulated the evolution of phenological cueing strategies in multiple environments, using historic climate data from 78 locations in North America and Hawaii to capture features of climatic correlation structures in the real world. Organisms in our model evolved diverse strategies that were spatially autocorrelated across locations on a continental scale, showing that similar strategies tend to evolve in similar climates. Within locations, organisms often evolved a wide range of strategies that showed similar response phenotypes and fitness outcomes under historical conditions. However, these strategies responded differently to novel climatic conditions, with variable fitness consequences. Our model shows how the evolution of phenological cueing strategies can explain observed variation in phenological shifts and unexpected responses to climate change.

## Introduction

Recent years have seen increasing interest in the study of phenological shifts. While organisms around the world have generally shown a “global coherent fingerprint” of advancing phenology with climate change (Parmesan and Yohe 2003; Parmesan 2007; Thackeray et al. 2010), several studies also point to substantial unexplained variation in phenological shifts (Parmesan 2007; Thackeray et al. 2010; Pearse et al. 2017). This variation in responses to climate change is an important factor driving phenological mismatch and the disruption of species interactions (Parmesan 2006; Kharouba et al. 2018). It has become increasingly clear that understanding how organisms integrate multiple environmental cues will be necessary to anticipate phenological shifts (Forrest and Miller-Rushing 2010; Visser et al. 2010; Pau et al. 2011; Chmura et al. 2019).

Although several studies have suggested factors that correlate with variation in phenological shifts (e.g., Parmesan 2007; Thackeray et al. 2010), relatively few studies have examined mechanistic explanations for this variation (Chmura et al. 2019). For example, while taxonomic groupings are often strong predictors of phenological shifts (Parmesan 2007; Davis et al. 2010; Thackeray et al. 2010, 2016; Davies et al. 2013), the mechanisms behind these groupings remain idiosyncratic or unclear (e.g., Parmesan 2007; Thackeray et al. 2010). The situation is similar with other proposed explanatory factors. Chmura et al. (2019) reviewed nine factors that have been suggested to structure variation in phenological shifts (including latitude, elevation, habitat, trophic level, life history, specialization, seasonal timing, thermoregulation and generation time), and concluded that most studies either do not suggest specific underlying mechanisms or do not evaluate alternative mechanistic hypotheses.

Our current study builds on previous modeling studies that explored phenological cueing strategies in a general context. These studies represent a progression from single-cue to multi-cue models, and towards more realistic environmental conditions. For example, Reed *et al*. (2010) used an individual-based model to examine plastic responses to simulated variation in a single cue and found that plasticity buffered fitness from environmental variation if the cue provided reliable information about environmental conditions, but had the opposite effect when the correlations between cues and conditions were weakened or when environmental variability was high. McNamara *et al*. (2011) developed a general analytical model based on a regression and correlation framework to explore the relationship between cues and optimal phenological timing under changing environments, and showed that environmental changes can affect the information value of cues in complex ways; as a result, they suggest that multiple cues could provide more robust predictive power than single cues. Chevin and Lande (2015) developed a multi-cue model to evaluate the evolution of multiple reaction norms in response to simulated environmental variation that included multiple correlated but fluctuating cues. This work showed that singular reaction norms can evolve to show plasticity that appears maladaptive when evaluated outside the multi-cue context, due to the correlated nature of environmental cues.

Here we present a generalized model that demonstrates how the evolution of integrated multi-trait cueing strategies can yield variable phenological responses to climate change. This model advances key themes established in previous studies by allowing cue integration strategies to evolve in the context of more complex, real-world environmental conditions. While previous modeling studies show that optimal multi-cue integration strategies depend on correlations among environmental cues (e.g. Chevin and Lande 2015), those results were based on simulated environments with known correlations. Our model aims to examine general mechanisms that emerge when organisms evolve to use the predictive information within real-world climatic data from different locations. Real-world climatic data are characterized by complex correlations among variables, and we assume that this correlation structure varies across locations, with relatively similar properties in nearby locations, and increasingly different properties in distant locations. Specifically, we examine how phenological cueing strategies could evolve to use correlations among climate variables to anticipate future events, and how these evolved strategies could contribute to observed phenological variation when historical correlations among climatic variables are disrupted.

We hypothesized that variation in cueing strategies could arise if organisms experiencing different environmental histories evolve different phenological strategies, caused by consistent differences in the reliability of predictive information provided by different kinds of environmental cues (Reed et al. 2010; McNamara et al. 2011; Chevin and Lande 2015). If the evolution of phenological cueing strategies was shaped by past environmental conditions in predictable ways, we expected that similar phenological strategies would evolve when organisms experienced similar historical climates. Conversely, variation among evolved strategies could persist under the same historical climate if different strategies were able to yield similar fitness outcomes. We further hypothesized that variation among evolved cueing strategies in their reliance on climatic and non-climatic cues could contribute to observed variation in phenological responses to climate change (Bonamour et al. 2019; Chmura et al. 2019).

## Model and methods

Our model simulates the evolution of a generalized, annual, asexually-reproducing organism in a simplified environment defined by daily maximum temperature, total daily precipitation, and day of the year (hereafter, temperature, precipitation and day). These conditions provide cues to anticipate future environmental conditions, and determine the fitness of individuals in the population (Fig. 1).

**Figure 1.**
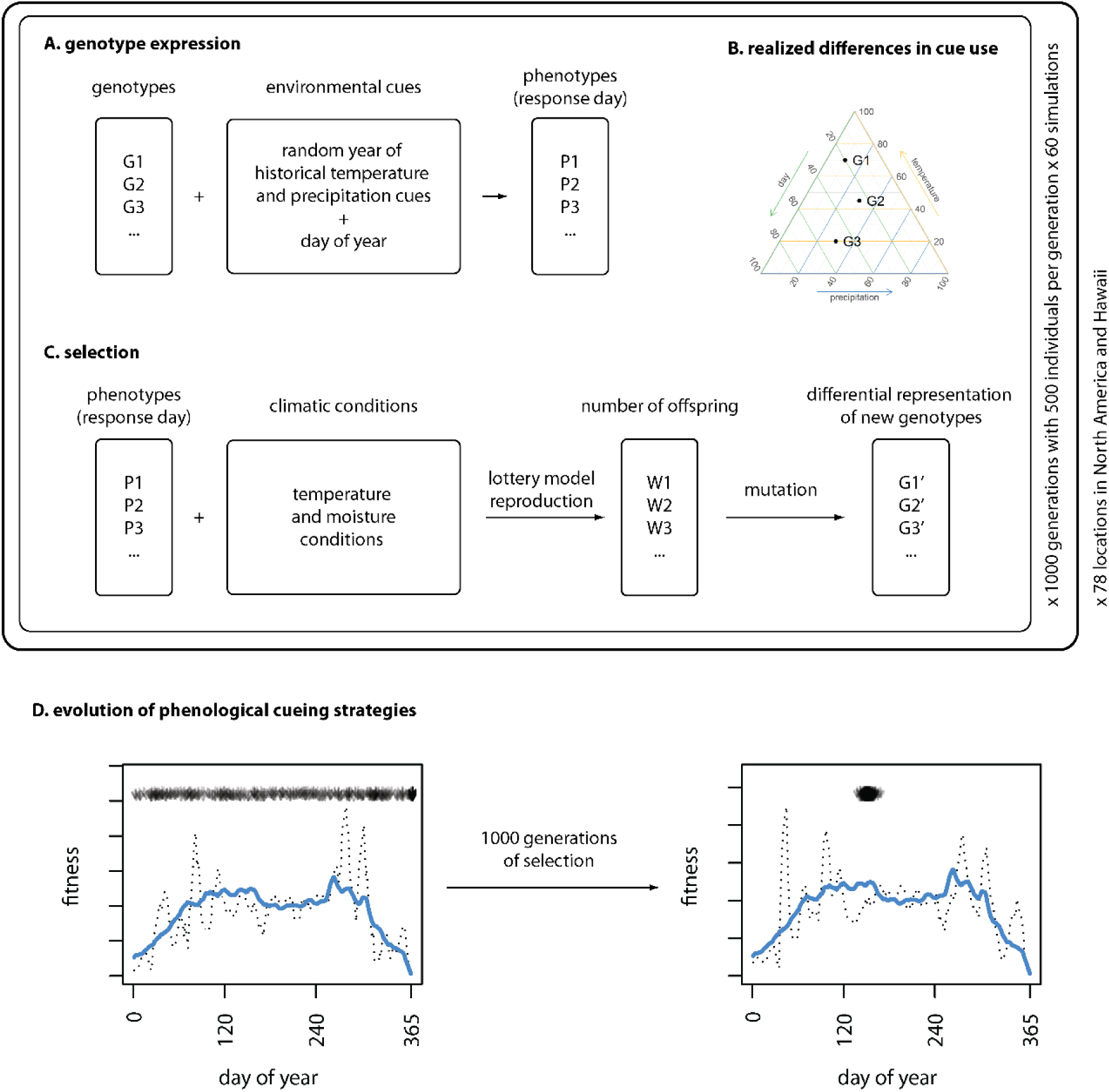
Schematic diagram of model. **A)** Genotypes combined with environmental cues (including cumulative annual daily temperature maximums, cumulative annual daily precipitation totals and day- of-year) result in expressed phenotypes (day of response). **B)** The trait effect (Τ), the proportional contribution of each trait to the response decision (a representation of the interaction between genotype and environment), can be expressed as a composition and presented on a ternary plot. **C)** The fitness of different phenotypes is determined by climatic (temperature and moisture) conditions during a 10-day window after the response threshold is crossed. A lottery model of reproduction determines the number of offspring produced by each individual, and mutation results in new genotypes for the next generation. **D)** Selection results in evolved phenological cueing strategies that anticipate favorable conditions and avoid unfavorable conditions. In this panel, the solid blue line represents the long-term expected fitness outcome for each day under historical conditions, while the dotted black lines represent the fitness outcomes for the first and last year of the simulation for the left and right panels respectively. The black arrows at the top of each panel represent the response day of each individual of the population. Initially the timing of phenological response is spread across the year, but after 1000 generations of selection, most of the population shows similar phenological timing. This example shows the results of one simulation using climatic data from Davis, CA, USA.

Our model combines three key features: 1) organisms combine multiple environmental cues using a weighted sum, 2) organisms make a phenological decision in response to a threshold of this weighted sum and 3) organismal sensitivity to each environmental cue is an evolved trait. Each of these features has been described across a wide range of organisms in nature (Gu et al. 2008; Wilczek et al. 2010; Burghardt et al. 2014; Seeholzer et al. 2018).

We implemented the simulation model and all analyses in R (R Core Team 2019).

### Cue integration

In our model, the set of environmental cues *E* is composed of cumulative annual daily maximum temperature (*γ*_*temp*_), cumulative annual daily precipitation (*γ_precip_*), and day-of-year (*γ_day_*):

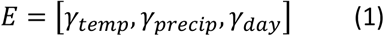

The set of environmental cues begins to accumulate on the first day of each year, and the cues change each day in each year of each location based on historical climatic data (we omit daily, yearly and location subscripts for simplicity in this notation, see *Environmental data* below). The use of cumulative annual temperature and precipitation assumes that organisms are aware of and can be influenced by past environmental conditions, consistent with degree-day models of development and phenology. Day-of-year provides a proxy for a consistent and non-climatic environmental cue, assuming that organisms are able to infer the day of the year (e.g., from photoperiod) with equal accuracy across all locations. Although the amplitude of seasonal photoperiodic changes is larger at higher latitudes, this assumption is supported by studies showing that tropical species are able to detect extremely small changes in photoperiod near the equator (Hau et al. 1998; Dawson 2007). More fundamentally, this assumption allows us to conservatively infer the relative information content of a climatically invariant cue across multiple locations, separate from the effect of increasing photoperiodic amplitude at higher latitudes. Using actual cumulative photoperiod produced qualitatively similar results (e.g., Fig. S1).

Each individual in our model has a genotype (G) defined by three traits (τ), which reflect its phenological sensitivity (sensu Chmura et al. 2019) to the three environmental cues:

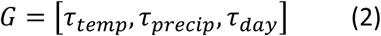

Each day of the simulation, each individual combines its cues and genotype into a weighted sum, which represents the response sum (S):

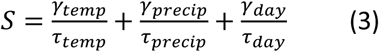

On the first day of the year when this sum exceeds the response threshold S≥1, the organism makes an irreversible phenological decision in anticipation of future fitness conditions. The genotype G thus represents the inverse weights of our weighted sum. We use 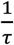 as the weights for the response sum for interpretability and consistency; genotypic traits are represented in the same units as the cue itself, and trait values indicate the critical cue value that would trigger a phenological response in the hypothetical absence of other cues. This also means that fixed increases or decreases to trait have the same effect regardless of trait value (*e.g.* increasing τ*_day_* by 1 means that in the absence of other cues the organism would respond one day later, regardless of whether τ*_day_* was previously 1 or 100). As a consequence, large trait values correspond to low sensitivity, and low trait values correspond to high sensitivity. Additive models of cue integration like this have been described in many organisms (Ernst and Banks 2002; Gu et al. 2008; Seeholzer et al. 2018), and similar assumptions have been applied in previous models (e.g., Jong 1990; Scheiner 1993; Lande 2009; Chevin and Lande 2015). While many organisms are likely to use more complex phenological cueing strategies across their life history (i.e., using multiple cues sequentially, as with chilling requirements for germination), additive models of cue integration provide a simple, commonly used, and plausible representation of how multiple cues are combined to form complex phenological cueing strategies.

### Fitness

Individuals reproduce at a rate proportional to the sum of the daily fitness they accrue over a fixed window starting one day after exceeding their response threshold. The fitness gained on any given day is the product of two skew-normal function outputs – one based on temperature, the other on moisture (see Eq. 4). These two fitness functions are combined to yield a 2-dimensional fitness surface akin to a quantitative version of a 2-dimensional Hutchinsonian niche (e.g., Fig. S2, Hutchinson 1957). Our model assumes that these two fitness factors interact multiplicatively rather than additively, so that favorable conditions in both dimensions are non-substitutable requirements for fitness, consistent with the Hutchinsonian niche concept. We used a skew-normal distribution because the thermal performance curves of ectotherms are generally asymmetrical, where fitness increases gradually as temperature increases towards the optimum, and then declines sharply above the optimum (Huey and Stevenson 1979; Sinclair et al. 2016). For simplicity, we used the same skew normal functional form (with a skew parameter of −10) for both temperature and moisture, though this model showed qualitatively similar results with alternative fitness functions (see *Sensitivity analyses*). Environmental moisture (*m*) was calculated based on daily precipitation totals (*p*) using a formula that includes a proportional retention constant (α) to represent the partial retention of moisture in the surrounding environment over time, as well as the input of new precipitation each day (p) (Eq. 4).

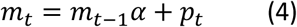

We set the retention constant to 0.8 in our simulations (but see *Sensitivity analyses*). At its limits, α = 0 represents daily precipitation, and α = 1 represents cumulative annual precipitation. We use α = 0.8 to reflect the assumption that organismal activity typically depends on moisture retained in the environment rather than daily precipitation. In contrast, cumulative annual precipitation was used in the cue integration model to reflect the assumption that organisms are aware of accumulated environmental information throughout the year. Changing the retention constant for environmental moisture produced qualitatively similar results, even when α = 0.

Temperature and moisture performance functions were parameterized separately for each location, such that the peak for each occurred at the 90^th^ percentile of all daily observations for a given location, and each function had a value that was 10% of the peak when the cue was at the 10^th^ percentile of all daily observations. This parameterization assumes that potential fitness values are maximized under relatively warm and moist conditions at each location, though this approach applies equally well to locations that are not characterized by these combined conditions because we simulate reproduction using a lottery model based on relative, realized fitness. Parameterizing by location without assuming performance constraints across sites (e.g., a universal minimum or maximum temperature across all locations) allows the interpretation of spatial patterns in evolved cue use without confounding differences in performance curves. This approach assumes that organisms are locally adapted to climatic conditions in a comparable way, so that evolved differences between locations are likely to be conservative, compared with a model in which universal constraints affect locations differently. To evaluate the robustness of observed results, we tested this model at a range of alternative fitness parameterizations, including different optimal quantiles, and observed qualitatively identical results (see *Sensitivity analyses*).

The raw fitness of each individual (W_i_) was calculated as the sum of these daily fitness payoffs over a 10- day window beginning one day after the response sum exceeded the response threshold (*e.g.*, Fig. S3). These raw fitness values combine the products of two probability densities and provide a relative measure of fitness; they have inherently small values and are not scaled to reflect expected numbers of offspring. Our model assumes a relatively short fitness accumulation window each year, where this window could represent any period when environmental conditions have a strong effect on fitness in the life history of our model organism. For example, it could represent the entire active, non-diapause phase of an organism’s life history or a short period of establishment in a longer life history (i.e., a seasonal window of opportunity sensu Yang and Cenzer 2020); for simplicity, we assume that environmental conditions do not have fitness effects outside of this window. Varying the duration of this window produced qualitatively similar results (see *Sensitivity analyses*). We use the sum of daily fitness payoffs to represent systems where the fitness benefits of favorable conditions accumulate over a window of time; while this is likely to represent some systems well, it would not adequately represent systems where daily fitness effects are multiplicative (e.g., a system in which a single extreme frost event has the potential to persistently damage flowers). We set this fitness window to begin one day after the response threshold is exceeded in order to simulate a delayed developmental process that required minimal anticipatory forecasting. Functionally, this one-day lag reduced the overall correlation between observed cues and experienced conditions. Increasing this lag further makes it more difficult for strategies to anticipate future conditions, but did not qualitatively change the behavior of this model (see *Sensitivity analyses*). All organisms in this model were constrained to have annual life histories with one generation per year; organisms that did not respond by the end of the year received zero fitness. This constraint prevented the evolution of multiple generations per year or multi-annual life histories, allowing us to focus on the seasonal phenology of relatively short-lived, annual organisms.

Individuals reproduced asexually with mutation (see *Heritability and mutation*), with population size held constant and expected realized fitness of each individual proportional to its calculated relative fitness. Reproduction was implemented as a lottery model to incorporate competition and allow for demographic stochasticity. With a constant population size each generation, individuals compete for representation in the next generation, with their probability of representation proportional to their raw fitness value (W_i_). For each evolved strategy (genotype) in the final generation, we calculated the geometric mean of its raw fitness across all years of environmental conditions. This fitness was proportional to its expected long-term relative fitness in that environment.

### Heritability and mutation

Offspring were given the same genotypes as their parent, modified by mutation. We modeled mutation by adding small random numbers (drawn from a normal distribution with mean 0 and a small standard deviation) to the parental traits. We set the standard deviation of mutation for each trait to be 0.5 percent of the overall cue range in order to produce mutation distributions with the same expected effect size in each location. In the case of the day cue, we used 360 as the maximum, leading to a standard deviation of 1.8 for mutation rate of the day trait in all locations. We assumed that each trait mutated for each individual in order to increase the overall rate of simulated evolution and improve computational efficiency.

### Environmental data

All available years of daily maximum temperature (degrees Celsius) and daily precipitation (mm rainfall equivalent) data were obtained from the NOAA Climate Data Online portal (Climate Data Online 2018) for 82 locations in North America and Hawaii. Locations were chosen to ensure spatial representation across the range of available data. After filtering for data quality and imputing missing daily values (see Appendix, *“Environmental data”*), we arrived at a climatic dataset of daily maximum temperatures and daily precipitation for 78 locations, with an average duration of 98 years (SD = 18.9 years, IQR = 114-84 years) (see Supporting Data Table 1).

To ensure that cue values were always non-negative, the temperatures for each location were shifted so that the minimum transformed temperature for that location was zero. Day of the year was represented as an integer value reflecting the number of days since January 1 of each year inclusive. The 366^th^ day was truncated from leap years in the dataset.

### Initialization and execution

For each location, we ran 60 simulations with the same parameter set. Each simulation included 1000 years (i.e., 1000 generations) of climatic data independently resampled from the historic dataset with replacement; as a result, the climatic history for each simulation was different, but drawn from the same historical climate distribution for that location (Fig. 1 and Supporting Table 1). Each simulation maintained a population of 500 individual organisms with individual genotypes. In the initial generation of each simulation, each individual was assigned uniform random trait values between 0 and 4 times the maximum cue value in that location (or 360 in the case of the day cue). This resulted in an initial population of individuals with a broad range of trait values (e.g., Figs. 1D and S4). Each simulation proceeded with the expression of a phenotype, the accumulation of resulting fitness payoffs, differential reproduction in a lottery model, and mutation in each generation. Thus, each simulation reflects a unique realization of the climatic history from a given location, with a randomly generated initial population. These simulations could be interpreted as replicates in an evolutionary experiment with systematic differences in the climatic means of locations, and stochastic variation in the specific sequence of climatic years and the specific genotypes of the initial population. Alternatively, each simulation could be interpreted as separate species that experience the same climate regime, and have the same temperature and moisture requirements.

### Assessing realized relative cue use

Trait values represent cue sensitivity; in our model, these can be interpreted as threshold values that would trigger a phenological response in the absence of other cues. Thus, the same trait values produce different behavior in different locations, depending on the environment. In order to compare strategies across locations, we define the “trait effect” (Τ) as a metric of proportional cue use. Each trait effect is a value from 0 to 1 that quantifies a strategy’s realized reliance on a given cue in a way that is comparable across locations. Specifically, this metric represents the proportion of the response sum *S* that is contributed by each 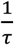 term of Eq. 3 on the day the response threshold is exceeded. Together, the trait effects of all three cues form a mathematical composition (here, a vector that sums 1) that represents realized cueing strategies in a way that is comparable across locations. Thus, we calculate mean strategies within simulations and locations using Aitchison compositional means (Boogaart and Tolosana-Delgado 2013), and plot these compositional means on ternary plots to show the three components of each strategy.

### Climate change scenarios

We examined how the individuals from the final generation of each simulation performed in novel climate regimes using two simple climate change scenarios. In the “*shift*” scenario, we advanced the historic temperature and precipitation regime in each year by 5 days. In the “*warming*” scenario, we increased all daily temperatures by 3 degrees, and left the precipitation regime unchanged. These two scenarios are not intended to represent detailed or realistic climate change scenarios; instead they reflect exposure to novel climates using a simple and systematic erosion of the key correlations in historical climatic data. In the *shift* scenario, the historical correlations between climatic conditions (temperature and moisture regime) and day-of-year are weakened, but the key correlation between the temperature and moisture regimes is maintained. Because of this, the seasonal fitness landscape (the timeline of potential daily fitness payoffs) in any given year remains unchanged, except that it is advanced by 5 days. In contrast, the *warming* scenario weakens historical correlations with temperature relative to the other two environmental cues, and also fundamentally changes the seasonal fitness landscape in any given year. Thus, while these two novel climate scenarios both represent important departures from the historical climate regime, the warming scenario presents a more profound departure from historical seasonal fitness landscapes. In both scenarios, we calculated the response day and fitness that would have been realized for each individual of our final populations in each unique year of the modified climate regime for each of 30 simulations. This allowed us to assess how climate change affected the phenotype and fitness consequences of each genotype that evolved under historical conditions.

For both climate change scenarios, we assessed correlations between each historic trait effect (Τ) and the change in response timing, and between each trait effect and the change in geometric mean fitness for each evolved genotype. We used linear mixed models with location as a random factor, allowing intercepts and slopes to vary. For these analyses, we report effect sizes (β) as the slope coefficient of each fixed explanatory factor; in these analyses, the effect size represents the expected change in response day or geometric mean fitness with a one unit change in the trait effect.

### Sensitivity analyses

We tested several model structures, cues, and parameter values to assess the robustness of our results (see Appendix, *“Sensitivity Analyses*”).

## Results

In many simulations, populations evolved to a region of successful trait combinations relatively quickly, with selection, mutation and drift leading to gradual shifts in the average population genotype, as well as the branching and pruning of lineages over time (e.g., Fig. S4). Some simulations experienced large shifts in trait use over time, often with concurrent changes ramifying across multiple traits. The individuals in the final generation typically emerged from the dominant evolved lineage, and showed similar combinations of traits. Thus, individual variation within each simulation was well-represented by the mean strategy for that simulation.

### Variation within locations

Mean evolved strategies often showed considerable variation between simulations, within locations. This variation in strategies can be observed in ternary plots (Figs. 2A-D and S5) and mapped to locations (Fig. S6). While some locations evolved tight clusters of similar strategies, most locations show a broad range of strategies using different sets of cues. These diverse strategies often showed geometric mean fitnesses that were similar to the most-fit mean genotype across all simulations (Figs. 2A-D S5, and S7). This occurred because most locations were characterized by high-performance fitness volumes that spanned a wide range of trait values, rather than a single clear optimal strategy. In 3-dimensional trait space, these high-performance fitness volumes resembled the hull of a boat or layers in a quartered onion (Figs. 2E-H and S8E-H), reflecting a wide range of evolved cueing strategies with similarly high geometric mean fitnesses (Figs. S7 and S9). These broad regions of trait space yielded similar fitness outcomes because they produced similar phenological behavior under historical conditions (Figs. 2I-L and S8I-L).

**Figure 2.**
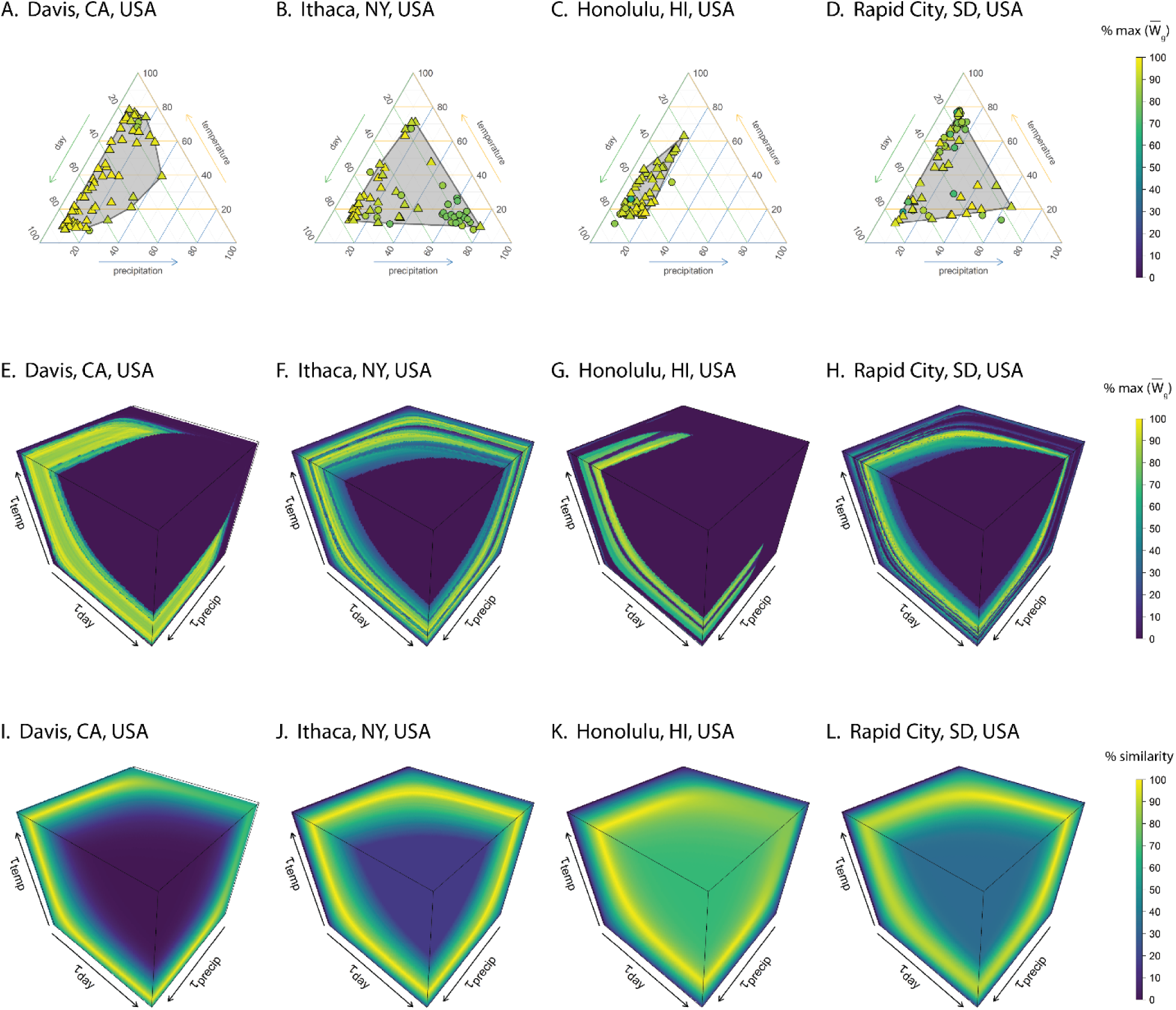
**A-D)** Ternary plots illustrate proportional cue use at the time of response for four selected locations. Each point represents the mean strategy at the end of one simulation; each strategy is represented as a composition of the trait effects (Τ) in percentages, which represent relative cue use. Point color reflects geometric mean fitness of genotypes across all years of the climate as a percent of the maximum observed geometric mean fitness 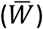 for each location. Simulations within 10% of the maximum observed geometric mean fitness in each location are represented as triangles and included in a gray convex hull. All other points are represented as circles. Ternary plots for all 78 locations are presented in Fig. S5, locations are described in Appendix Table 1. **E-H)** Geometric fitness in the 3- dimensional trait-space of our organisms, with each dimension representing phenological sensitivity to a different cue (where low trait values mean high sensitivity). For each location, the yellow region represents strategies that were at or near the highest observed fitness. This region generally spans a wide range of trait values, reflecting the breadth potentially successful trait combinations. These plots show that diverse genotypes can produce similarly high fitness phenotypes. To generate these plots, we evaluated a 100×100×100 grid spanning trait values ranging from the 10^th^ through 90^th^ percentiles of observed cues in each location for fitness and response day in each recorded year of climate. **I-J)** Response similarity is plotted for the same trait ranges in each location. Response similarity is a metric that quantifies the proportional similarity of phenological responses for each trait combination (genotype) compared with the phenological responses of the trait combination with the maximum geometric mean fitness across all available years. We calculate the response similarity as one minus the proportional response dissimilarity, which was defined as the mean distance (in days) between the response day of each genotype compared with that of the most fit genotype, divided by the greatest distance in each year. Bands of high similarity span most of the traitspace, demonstrating that many different combinations of traits can lead to similar response phenologies. Comparisons with panels E-H show that regions of high response similarity generally overlap with regions of high fitness, illustrating that the observed similarity in fitness between diverse strategies is generally due to the expression of similar phenological phenotypes, rather than alternative phenotypes with equivalent fitness.

### Variation between locations

We found considerable spatial variation in the mean evolved strategies across locations. Spatial patterns in mean cue use are visible when mapped (Fig. 3A), and showed strong positive autocorrelation on a continental scale (Fig. 3B-D). This result indicates that similar mean strategies evolved under similar climates, suggesting a degree of underlying predictability in the evolution of phenological cueing strategies, despite the variability of evolved strategies among simulations within each location.

**Figure 3.**
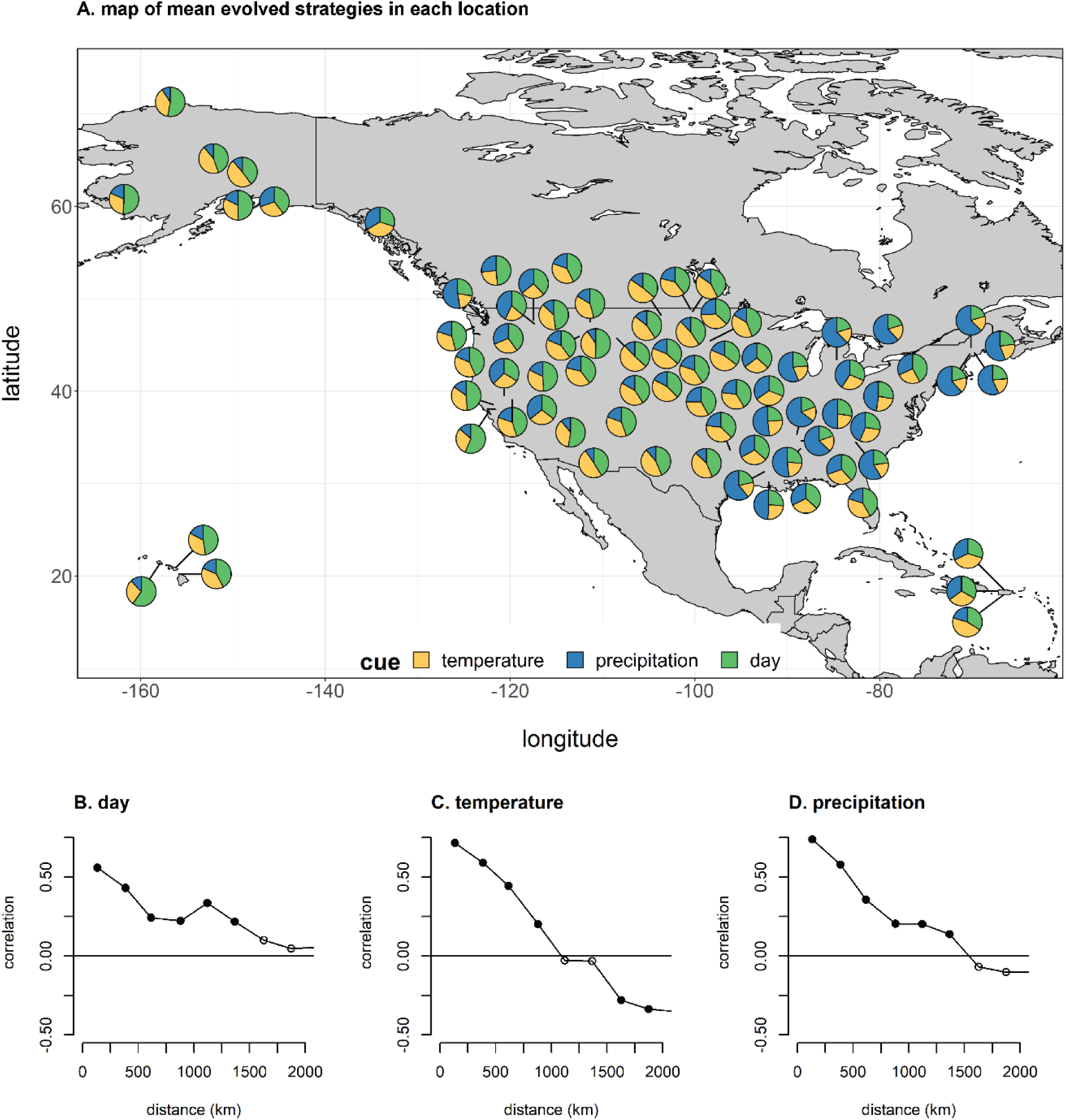
Evolved strategies show spatial autocorrelation in relative cue use (the trait effect, Τ); similar strategies evolve in nearby locations with similar climates, and different strategies evolve in distant locations with different climates. **A)** A map representing the mean evolved strategy for each of 78 locations across all years. Evolved strategies show significant positive spatial autocorrelation (Moran’s I) in reliance on **B)** day, **C)** temperature, and **D)** precipitation cues up to at least 1000 km. Filled circles are significantly correlated, open circles are not.

We analyzed several climatic and location variables as potential correlates of evolved mean cue use at each location (Appendix *“Analysis of explanatory factors”*, Fig. S10 and Fig. S11). While several factors emerged as potentially meaningful predictors of phenological cue use in this analysis, most of the variation in cue use was unexplained even in models that combined all 17 factors (Τ_day_: marginal R^2^=0.102; Τ_temp_: marginal R^2^=0.110; Τ_precip_: marginal R^2^=0.308).

### Responses to climate change scenarios

In the *shift* climate change scenario, populations generally advanced their mean phenology, but showed highly variable changes in their realized fitness when comparing between (Fig. 4A) and within locations (Fig. 5A,C,E,G). These effects were non-random; as expected, organisms that relied more on day cues were less likely to advance their phenology on pace with the changed climate (β=4.6 days per unit Τ, SE=0.02, p<0.0001), while organisms that relied more on temperature or precipitation cues were more likely to advance their phenology (temperature, β=-2.5 days per unit Τ, SE=0.19, p<0.0001; precipitation, β=-2.2 days per unit Τ, SE=0.1, p<0.0001). Because the seasonal fitness landscape retained the same shape but advanced by 5 days in this scenario, organisms that relied more on day cues generally showed a weak pattern of more negative fitness consequences (β=-.00023 units W_i_ per unit Τ, SE=0.00007, p=0.0009, Fig. 4A), while those that relied more heavily on temperature or precipitation cues showed weakly positive fitness consequences (temperature, β=0.00017 units W_i_ per unit Τ, SE=0.00005, p=0.002; precipitation, β=0.00009 units W_i_ per unit Τ, SE=0.00005, p=0.04). While most locations experienced a reduced mean fitness under the changed climate, some locations showed higher overall fitness (Fig. 4A). Simulations within locations showed similarly variable responses in both advancement and fitness (Fig. 5A,C,E,G). The behavior of individual genotypes within each location (Fig. S12A) shows how small changes in phenological response phenotype can drive large changes in mean fitness outcomes under the *shift* scenario.

**Figure 4.**
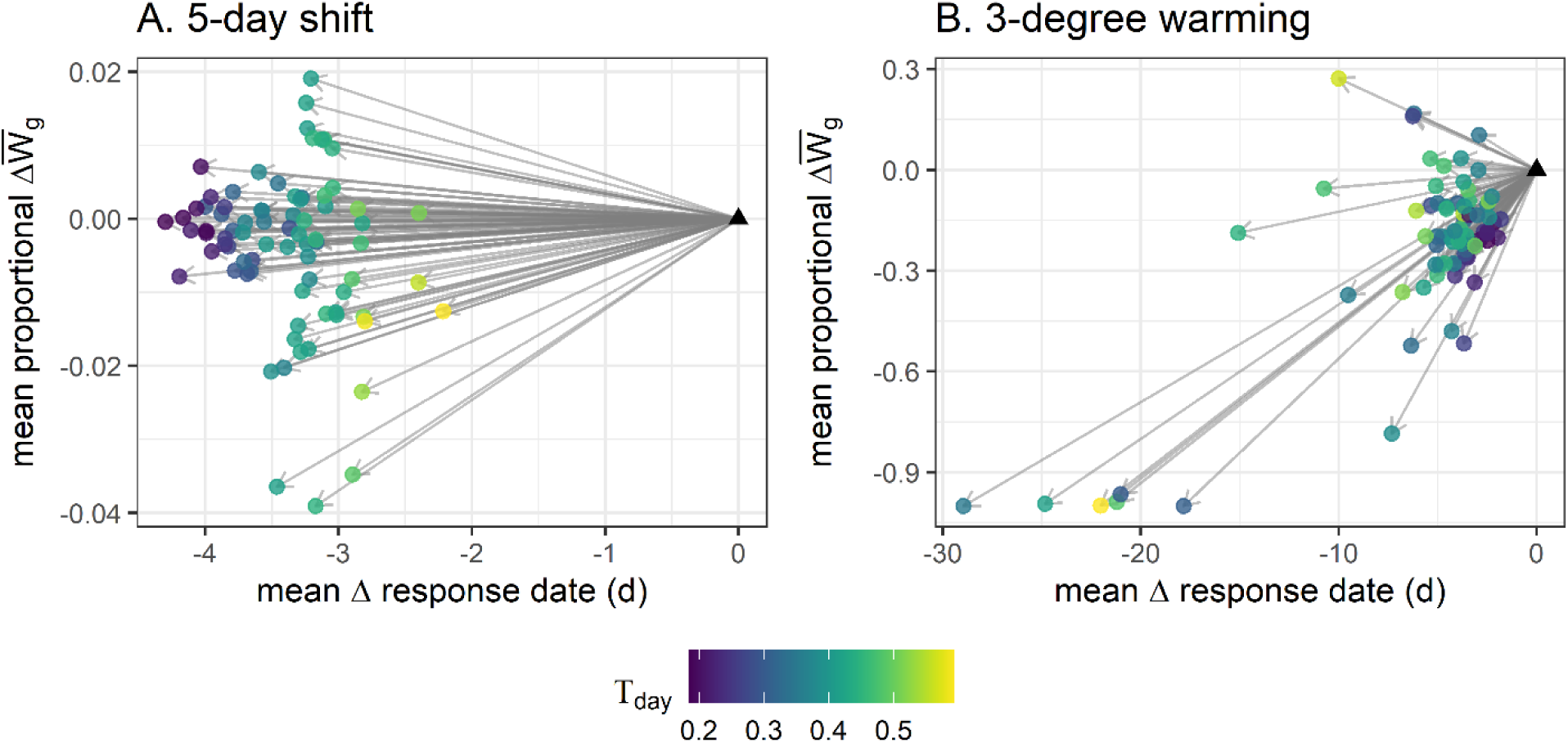
The variability of mean phenological responses to climate change scenarios and their fitness consequences across locations (see also Figure 5 for variation across simulations within locations). Changes in phenology and mean fitness **A)** under a *shift* scenario where both temperature and precipitation regimes advance by 5 days, and **B)** under a *warming* scenario where temperature are warmed by 3-degrees across the year. Under these two scenarios, organisms with different environmental histories generally respond earlier, but show variable degrees of advancement and highly variable fitness consequences. The position of each circle represents the mean change in the response date and the proportional change in geometric mean fitness averaged across all evolved genotypes of 30 simulations for each of 78 locations, relative to the historical climate in that location, represented by a black triangle. Thus, this figure shows the variability of phenological responses and fitness consequences to climate change across locations (see Fig. S12 for plots of phenological responses and fitness changes across all locations). The color of each circle represents the historical trait effect of day (Τ_day_), indicating the relative use of a climatically invariant cue.

**Figure 5.**
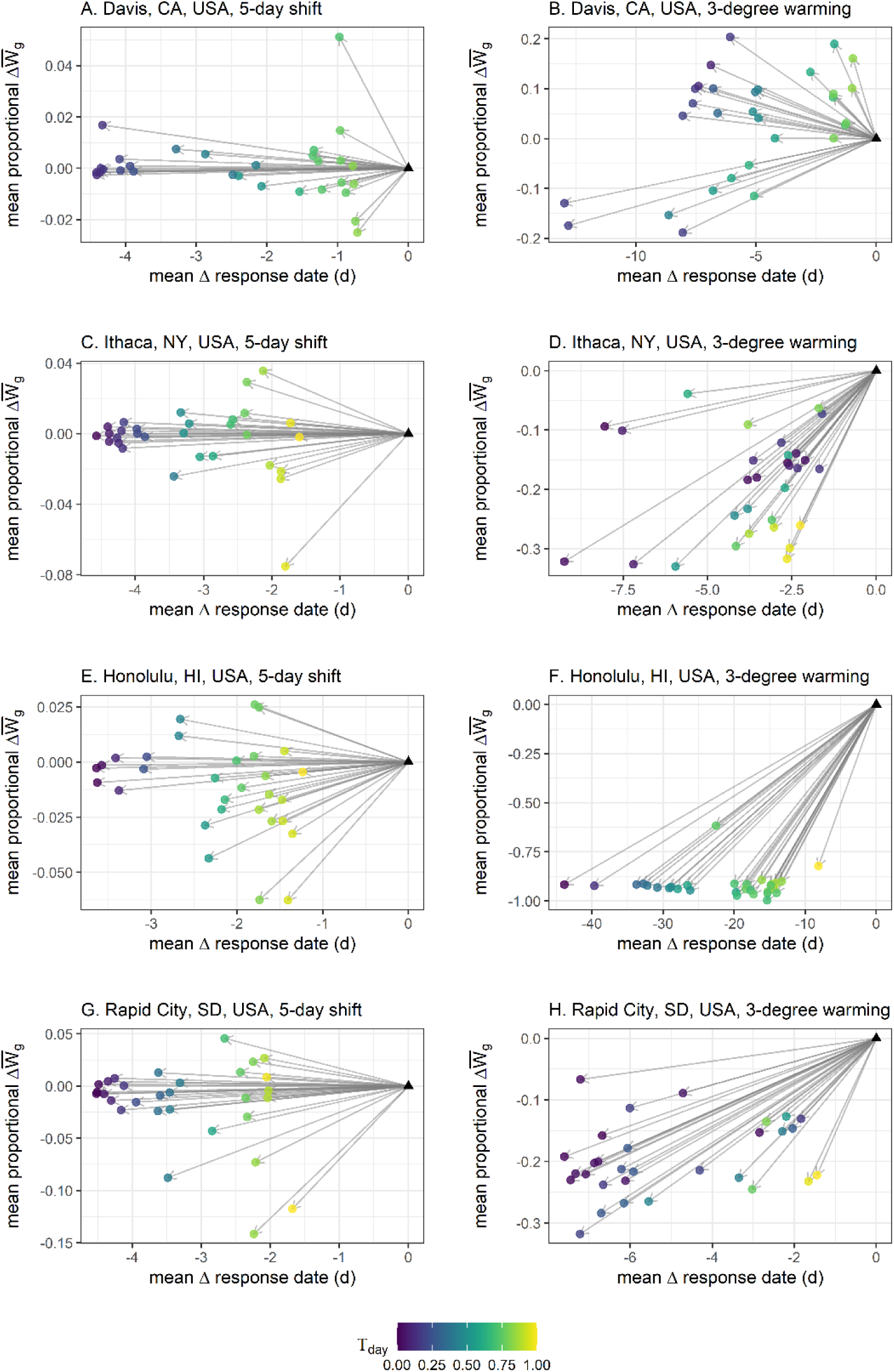
The variability of mean phenological responses to climate change scenarios and their fitness consequences across simulations within locations (see also Figure 4 for variation across locations). This figure is constructed as in Figure 4, but showing changes in phenology and mean fitness for each of 30 simulations from four example locations (rows) under both the shift scenarios (left column, panels **A**, **C**, **D**, and **G**) and the warming scenario (right column, panels **B**, **D**, **F**, and **H**). Each colored circular point represents the mean response of a single simulation relative to the historical condition, represented by a black triangle. The color of each circle represents the historical trait effect of day (Τ_day_), indicating the relative use of a climatically invariant cue.

Under the *warming* scenario, populations also advanced their mean phenology, both between (Fig. 4B) and within locations (Fig. 5B,D,F,H). Mean strategies with greater reliance on day or precipitation cues showed reduced phenological advancement (day, β=8.1 days per unit Τ, SE=1.2, p<0.0001; precipitation, β=7.7 days per unit Τ, SE=1.0, p<0.00001), while those that relied more on temperature showed greater phenological advancement (temperature, β=-15.4 days per unit Τ, SE=1.5, p<0.00001). This effect of day was more apparent in the *shift* scenario than the *warming* scenario when comparing across locations (Fig. 4A vs. 4B), but is apparent in both scenarios when comparing within locations (Fig. 5). Organisms with greater reliance on day and precipitation showed higher fitness in the *warming* scenario (day, β=0.0006 units W_i_ per unit Τ, SE=0.0002, p=0.0026; precipitation, β=0.0007 units W_i_ per unit Τ, SE=0.0003, p=0.01), while those that relied more on temperature showed reduced fitness (β=-0.001 units W_i_ per unit Τ, SE=0.0002, p<0.00001). Many locations showed larger and less predictable changes in mean fitness outcomes under the *warming* scenario than in the *shift* scenario (e.g., Figs. 4, 5 and S12).

### Sensitivity analyses

Our findings were qualitatively robust across a wide range of model variants using different cues (daily precipitation, daily temperature, photoperiod, quadratic measures of cues), fitness functions, and fitness window durations (see Appendix, *“Sensitivity analyses*”). Differences in historic dataset length did not explain a meaningful proportion of the climatic variation across locations (Appendix, *“Sensitivity to dataset length”*).

## Discussion

This model suggests two key findings. First, we see that the evolution of phenological cueing strategies was shaped by environmental history in broadly predictable ways across locations (Fig. 3), despite substantial variation in cueing strategies within locations (Figs. 2A-H and S5). Second, evolved cueing strategies showed highly variable responses to simulated climate change (Figs. 4 and 5).

### Predictability and variation in the evolution of phenological cueing strategies

The observation that similar mean strategies evolved in locations with similar climates likely reflects continental scale spatial patterns in the relative reliability of temperature, precipitation and day cues (Fig. 3). The spatial autocorrelation of evolved strategies indicates that evolution tended to produce similar phenological cueing strategies under similar environmental histories. Our model assumes that selection will favor cues based on their ability to predict future environmental conditions that are relevant to fitness – both the ability to consistently trigger a phenological response ahead of favorable conditions, and the ability to avoid triggering a phenological response ahead of unfavorable conditions. Thus, this result suggests that broad patterns in phenological cue use may be predictable based on the relative information content of different cues.

Across locations, we observed a broad and complex range of strategies evolving in response to real-world climatic data. In some locations, this resulted in strategies that relied heavily on climatic cues to track factorable climatic cues across year-to-year variation (e.g., Figs. 2 and S5). In other locations, this resulted in the evolution of bet-hedging strategies with a greater reliance on climatically invariant day-of-year cues (e.g., Figs. 2 and S5). While broad patterns of phenological cue use may be predictable based on the relative reliability of environmental cues in an organism’s evolutionary history, simple climatic or location variables were only marginally successful at characterizing the relevant differences between locations in our model, and the majority of observed variation in evolved cue use could not be explained by a model including all evaluated climatic and location variables (Appendix, *“Analysis of explanatory factors”*, Figs. S10 and S11). This likely reflects the fact that most of the *a priori* descriptive variables we used were too coarse, static or general to capture the aspects of climatic predictability that are most relevant to our model organisms. For example, many of these descriptive variables were metrics of annual climatic variability averaged across years, and such general descriptors likely failed to capture the specific predictability of cues in most relevant part of the season for our model organisms.

In addition to the variation we observed in mean cueing strategies between locations, we also observed substantial variation in evolved cueing strategies across simulations within locations (Figs. 2A-D and S5). This variation emerges because a wide range of trait value combinations (i.e., cueing strategies or genotypes, Fig. 2A-D) yield similar response phenotypes (e.g., Fig. 2I-J) with similar fitness outcomes (e.g., Fig. 2E-H). This is a fundamental consequence of multi-cue integration when there are correlations among climatic cues; when these conditions are met, changes in one component of a cueing strategy can often be compensated for through changes in another. This feature of cue integration can lead to a broad range of multi-cue strategies that can appear counterintuitive when singular cue responses are examined in isolation (c.f., Chevin and Lande 2015). This fundamental property of multi-cue integration predicts that organisms showing similar phenologies under historical climates could have widely divergent underlying phenological strategies that use different cues to different degrees. This prediction is consistent with the findings of empirical studies showing differences in cue use between interacting species that generally show phenological synchrony (e.g., Iler et al. 2013).

This fundamental consequence of cue integration is further complicated by non-additive interactions among traits in our model. This is partly due to our response threshold model, which creates inherent non-linearities in the relationships between traits and the phenotype. It also reflects the variable nature of real-world climatic dynamics across each year, which cause the effects of one trait to depend on the effects of the other traits in an individual’s genotypic background in complex and non-additive ways. As an extreme example, a trait that confers a very high sensitivity to one cue can nullify the effects of other cues, because even a small value of one cue will cause the organism to exceed its response threshold. More generally, the effects of any trait on both phenotype and fitness depend on the other traits in the organism’s strategy and the seasonal dynamics of its environment. These non-additive interactions between traits are akin to epistasis (Phillips 2008), and create the potential for a more diverse and complex range of cueing strategies with similar fitness outcomes in any given location (Fenster et al. 1997).

### Novel climates result in ecological surprises

Our second key finding is that phenological strategies which produced similar phenotypes under historical conditions showed strong phenotypic and fitness differences under simulated climate change (Figs. 4, 5 and S11). These effects depended on the degree to which our climate scenarios broke key correlations in the historic climate. In the *shift* scenario, temperature and precipitation regimes were advanced in unison, but the correlations between temperature and precipitation were unchanged. Thus, organisms that were more sensitive to climatic cues showed greater phenological advancement and more positive fitness consequences, while those that relied more heavily on climatically invariant day-of-year cues showed reduced advancement and more negative fitness consequences (Figs. 4, 5 and S11). This result is consistent with expectations about the costs of using invariant day-of-year (e.g., photoperiodic) cues under climate change (Coppack and Pulido 2004; Way and Montgomery 2015). However, in the *warming* scenario, greater reliance on the invariant day-of-year cue was generally favorable, while organisms that relied more on temperature unexpectedly showed reduced fitness (Figs. 4, 5 and S11). This result occurs because the *warming* scenario increased temperatures independently of precipitation, thus breaking historic correlations between temperature- and precipitation-based factors. Because fitness is a function of both temperature and precipitation in our model (Fig. S2), the warming scenario changed the seasonal fitness landscape in complex and novel ways. These changes to the seasonal fitness landscape made the fitness consequences of phenological advancement less predictable. As a result, many locations showed large and counterintuitive changes in mean fitness outcomes under the *warming* scenario (Figs. 4, 5 and S11).

A comparison of the *shift* and *warming* scenarios suggests some general insights. The specific ways in which these two scenarios differed had important consequences. The increased unpredictability of fitness responses under the *warming* scenario suggests that even a relatively small decoupling of the historical temperature and precipitation regimes could increase the likelihood and costs of maladaptive plasticity. These results are consistent with the hypothesis that organisms are more likely to show maladaptive and counterintuitive plasticity in environments that differ most from those in their evolutionary history (Ghalambor et al. 2007; Chevin et al. 2010; Reed et al. 2010; McNamara et al. 2011; Chevin and Lande 2015; Duputié et al. 2015). Thus, while intuition suggests that a greater reliance on climatic cues (as opposed to climatically invariant cues) would allow for more adaptively plastic responses to a changing climate, our findings suggest that this may not always be the case.

At the intersection of our two key findings, a wide range of strategies which show predictable and consistent behavior under historical conditions can show unpredictable and counterintuitive behavior in a novel environment. This is consistent with the idea that multi-cue phenological strategies create the potential for cryptic genetic variation to be expressed under climate change. Cryptic genetic variation is genetic variation that is not normally expressed, but which can yield phenotypic variation under changed conditions (Rutherford 2000; Gibson and Dworkin 2004; Gibson and Reed 2008; McGuigan and Sgrò 2009; Paaby and Rockman 2014). Cryptic genetic variation appears to be widespread in eukaryotes, and may be particularly characteristic of systems where response thresholds provide a mechanism of “genetic buffering” (Rutherford 2000). In the context of our model, the mechanisms that maintain genotypic variation while minimizing fitness differences under historic climate conditions could contribute to the maintenance of cryptic genetic variation, increasing the potential for ecological surprises under novel climates.

### Context and broader implications

Our model examines the evolution of multi-cue strategies and its implications for variation in phenological responses to climate change. Previous studies have identified important patterns of phenological shift in nature (e.g., Parmesan 2007; Thackeray et al. 2010), and examined the behavior of increasingly complex phenological cueing models under increasingly realistic simulated environments (e.g., Reed et al. 2010; McNamara et al. 2011; Chevin and Lande 2015). Our current study provides a complementary approach to examine how evolution and cue integration could affect patterns of variation in phenological shifts. We find that phenological cueing strategies that evolve in the context of real world climatic data show patterns of cue use that can be broadly understood in the context of cue reliability, consistent with previous modeling studies (Reed et al. 2010; McNamara et al. 2011). However, these evolved patterns of cue use can also show a great deal of complex and sometimes cryptic variability, consistent with our understanding of multi-cue integration from previous models (e.g., Chevin and Lande 2015). This variability in evolved cue use can lead to high variability in phenological responses to climate change, with phenotypic and fitness consequences that are increasingly difficult to predict under increasingly novel climate regimes. This result is consistent with expectations about the limits of adaptive plasticity in novel environments (e.g,, Ghalambor et al. 2007; Chevin et al. 2010; Duputié et al. 2015), and suggests that the many organisms may show increasingly counterintuitive responses to climate change.

Our model results suggest that we should expect to see substantial variation in phenological shifts, even if organisms experience similar environmental changes. Chmura et al. (2019) proposed a key distinction between organismal and environmental mechanisms of variation in phenological shift, with the former driven by differences among organisms in their sensitivity to cues, while the latter is driven by differences in the environmental change that different organisms experience. In this context, our model is focused on the evolutionary origins of variation in organismal mechanisms. In our climate change scenarios, we control and hold constant the environmental change that each population experiences. As a result, the substantial variation in phenological responses to climate change observed in our simulations is driven by differences in cueing strategy. The results of our model suggest that even when we limit the potential mechanisms of variation in phenological shifts, evolved differences in cueing strategies would contribute to a great deal of observed variation in phenological responses to climate change.

### Scope, aims and limitations

Our model was developed to explore general mechanisms for the variability of phenological shifts, and does not attempt to make quantitative predictions about the evolution of cueing strategies at specific locations for any specific organism. For example, the patterns of cue use shown on the map in Fig. 3A represent only one possible model outcome, generated under one set of model parameters and assumptions. In the absence of a specifically parameterized model, these results should not be interpreted as meaningful predictions for any given system. We present this figure as an example to illustrate a more general finding - that similar mean strategies tend to evolve in locations with similar climates, while different strategies tend evolve under different environmental histories. Unlike specific patterns of cue use in specific locations, this is a robust result that we see across a wide range of model parameters. Although more specific questions will require a more detailed models, we hope that this general theoretical framework will encourage more specific studies in the future.

### Future directions

Future studies could extend this model by increasing model complexity, or evaluating our general findings in specific systems. Potential extensions of this model include modeling organisms with alternative life histories, using a broader range of environmental cues, considering more complex cue integration mechanisms, allowing sexual recombination of traits to increase standing genetic variation, or allowing gene flow between locations. It would be particularly useful to study whether more complex cueing strategies could allow greater resilience or robustness in the face of climate change. However, our ability to apply models to make predictions relevant to specific systems is likely to be more limited by our current knowledge of key parameters in specific systems rather than our ability to develop more complex models. Future empirical and observational studies could build a groundwork for these studies by identifying key cues and cue integration mechanisms, and by documenting variation in phenological cueing strategies within and across populations.

While we used different locations to represent different environmental conditions in this model, the general findings of this model could also potentially be extended to consider other factors that structure the availability of environmental cues, such as microhabitats or life histories. Two organisms in the same location may experience very different environmental conditions, potentially structured by their microhabitat, life history, trophic position, body size, or other factors. For example, the general findings of our model could potentially be applied to observed differences in phenological shifts correlated with phylogenetic groups (e.g., Parmesan 2007; Davis et al. 2010; Thackeray et al. 2010; Davies et al. 2013). Parmesan (2007) speculated that the particularly strong and variable phenological shifts of amphibians could be due to their particular reliance on precipitation-associated cues. Similarly, Davis et al. (2010) hypothesized that phylogenetic patterns in flowering time shifts could be caused by differences in cue use, potentially reflecting differences in the reliability of different cues in the evolutionary histories of different taxa. The results of our model are consistent with these hypotheses, and the expectation that organisms exposed to different environments over their evolutionary history will evolve different phenological cueing strategies, with consequences for their phenological responses to climate change.

### Conclusion

The two key findings we report here are robust across a range of model parameters, and appear to be rooted in fundamental mechanisms of multi-cue integration and the complexity of real-world climatic correlations. This suggests that similar mechanisms could potentially occur in a wide range of systems (e.g., Beshers and Fewell 2001; Wilczek et al. 2010; Seeholzer et al. 2018; Chmura et al. 2019), and that examining the reliability of cues in an organism’s evolutionary history could provide useful a starting place for understanding current phenological cueing strategies. Understanding current phenological cueing strategies could potentially improve our ability to predict and respond to future phenological shifts. However, these results also suggest that the nature of cue integration may put fundamental limits on our ability to predict the responses and fitness outcomes of organisms living under novel climatic regimes.

---

**Figure S1.**
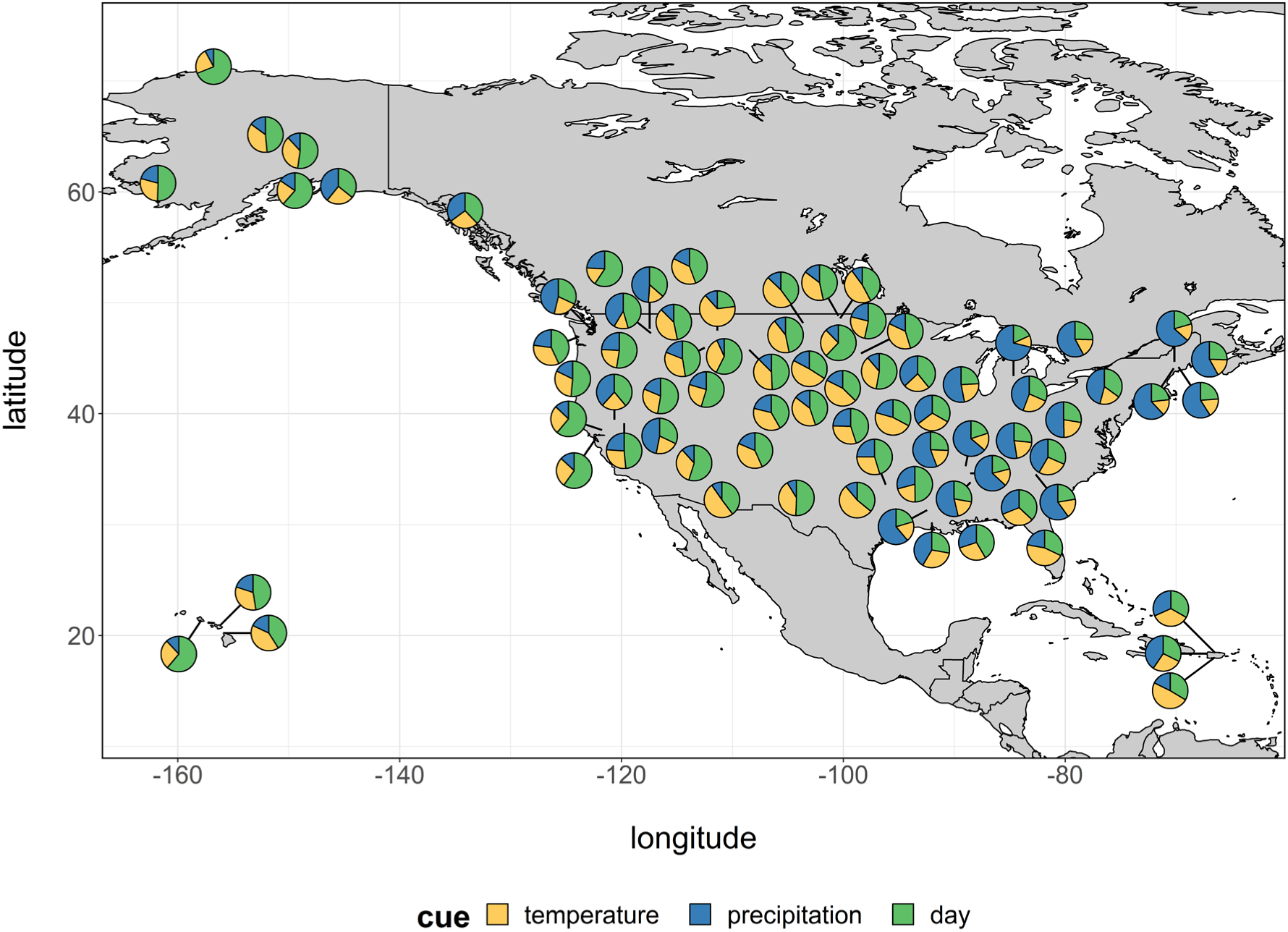
A map of mean phenological strategies in different locations under an alternative model formulation using cumulative photoperiod as the cue for day-of-year an a developmental baseline temperature of 0 C across all locations below which organisms are insensitive to thermal cues. As in the primary model formulation, similar strategies evolve in similar climates.

**Figure S2.**
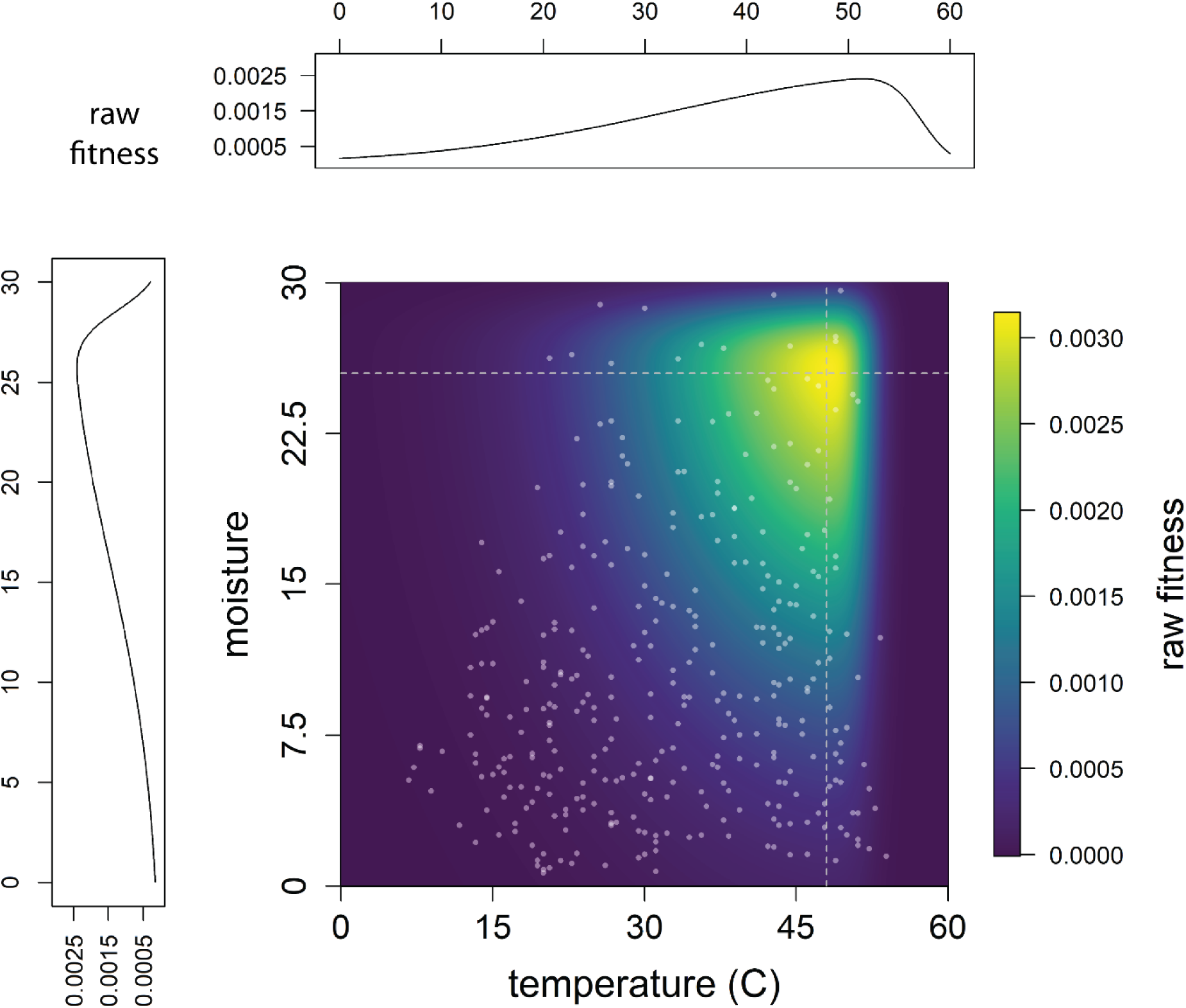
A visualization of the 2- dimensional fitness function and example seasonal fitness landscapes. Daily fitness function, parameterized for Ithaca, NY, USA showing the 2-dimensional skew-Gaussian shape; colors represent raw daily fitness. White points show daily measures from the first year of Ithaca data, 1893. Side panels show the 1-dimensional asymmetric fitness function associated with temperature and moisture at slices through the 2-dimensional surface, represented by the dashed lines.

**Figure S3.**
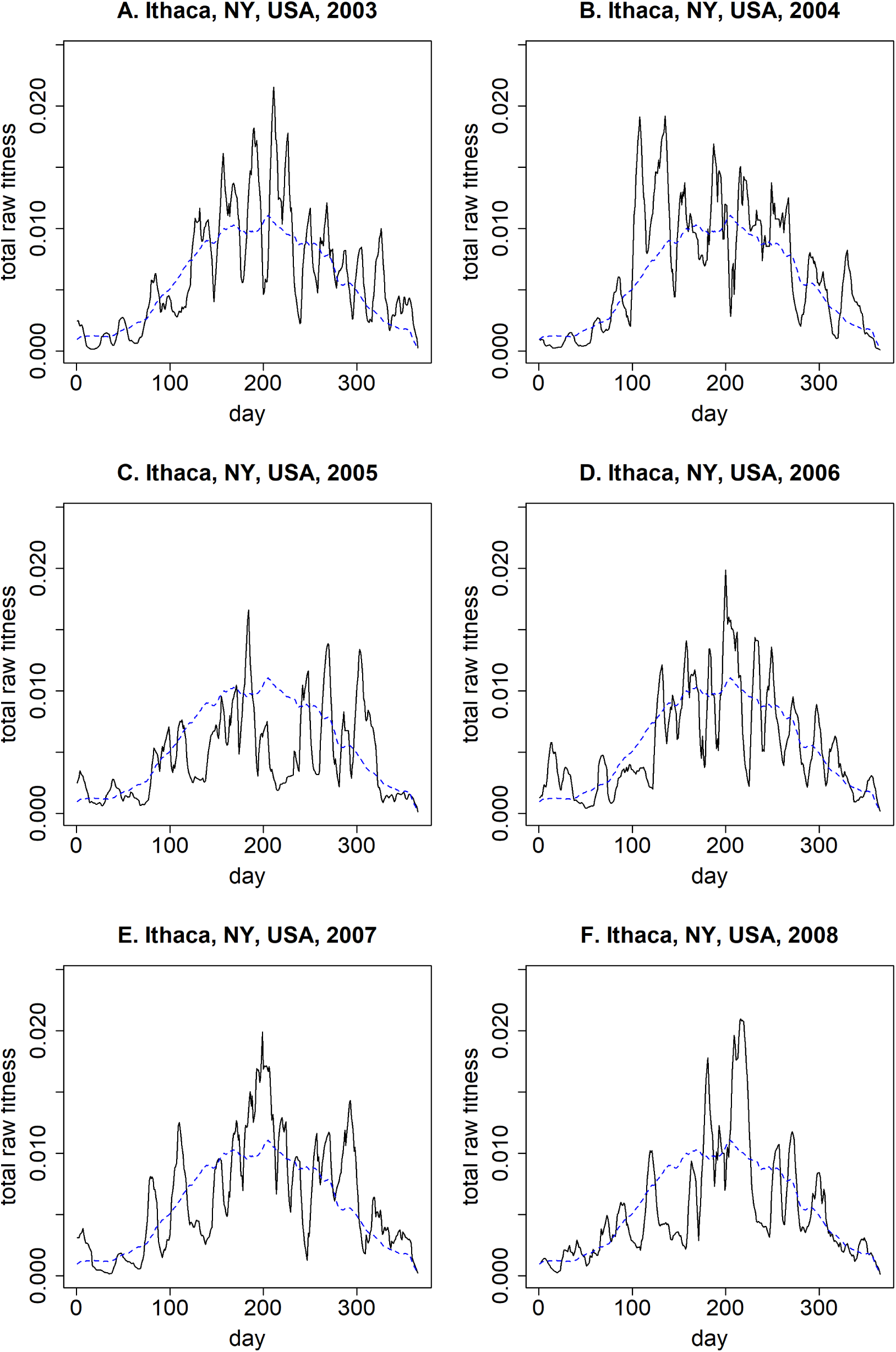
Examples of the seasonal fitness landscape for Ithaca, NY, USA in 2003-2008. The y-axis represents the total accumulated raw fitness payoff over a 10-day window as a function of phenological response timing (day-of-year). The blue dashed line represents the expected fitness landscape for any given year, based on the long-term mean.

**Figure S4.**
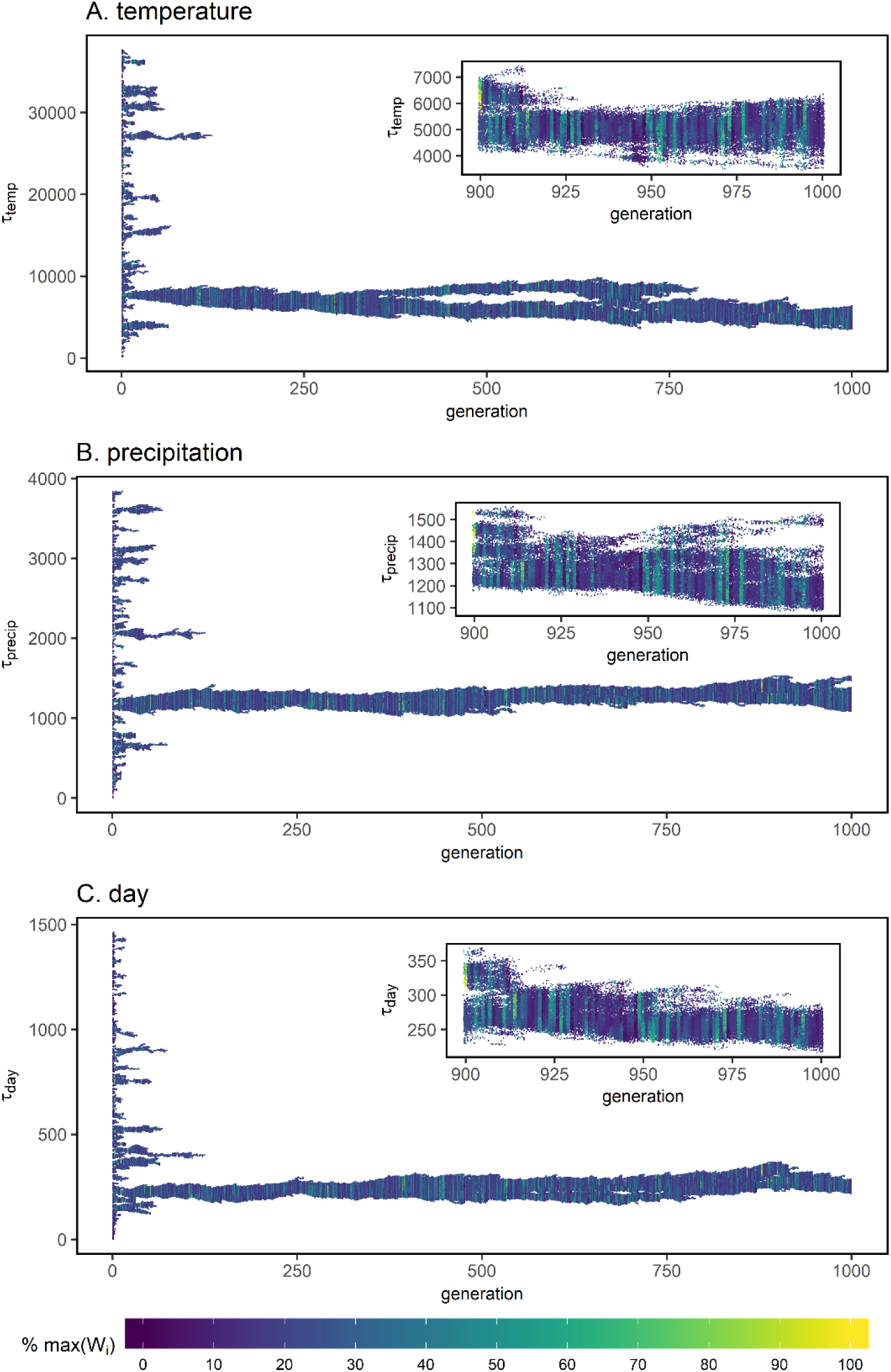
Temperature, precipitation and day traits evolve over 1000 generations in this representative simulation from Davis, CA, USA. Each simulation begins with initial values for each trait drawn from broad uniform distributions; selection drives the evolution of specific strategies. The inset figure shows an expanded view of trait evolution in the final 100 generations. Each point represents the trait value of one individual; point colors show raw individual fitness proportional to the maximum value in this simulation. Populations commonly experienced “good” and “bad” years, and often showed coordinated changes in trait values across the three cues.

**Figure S5.**
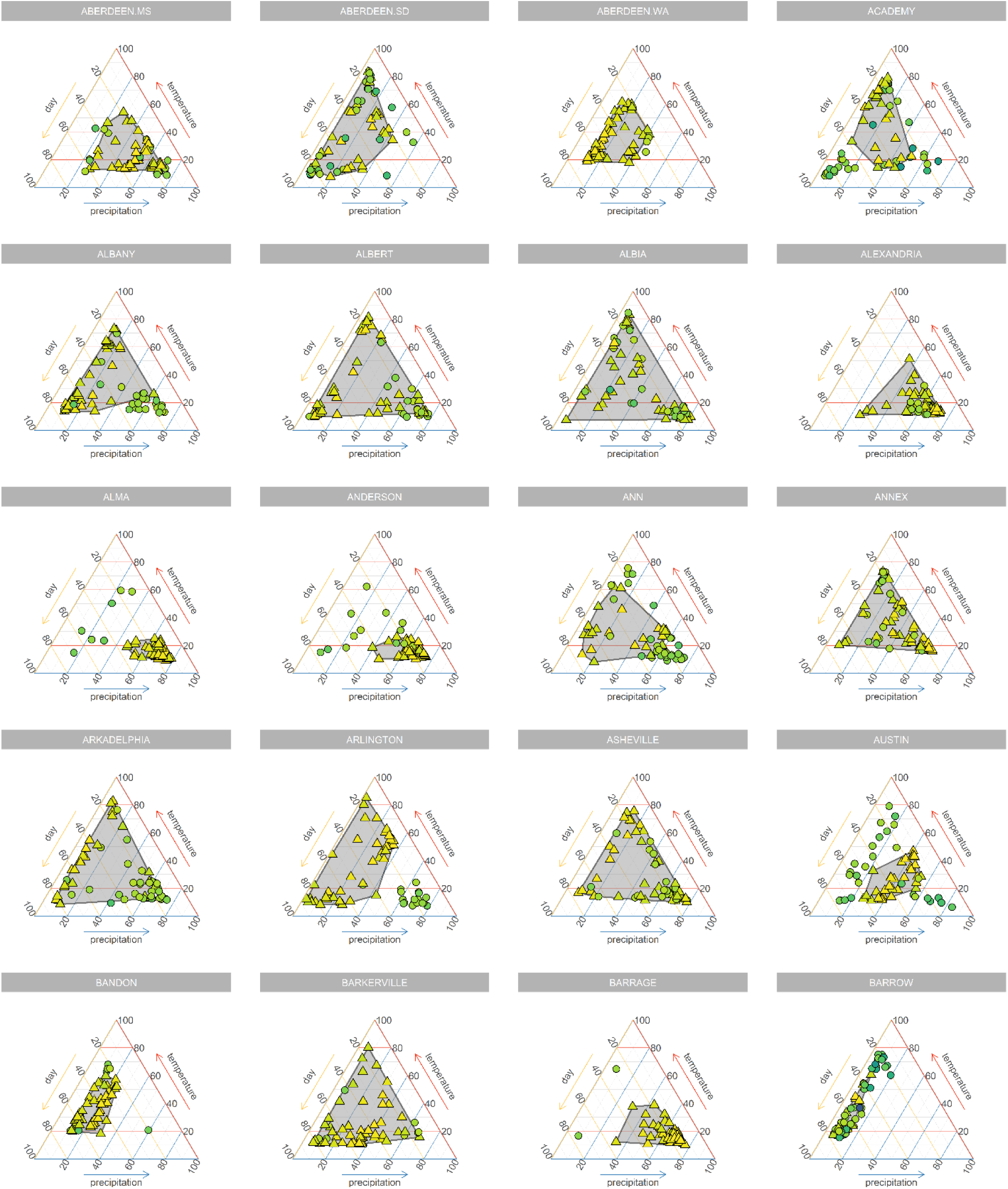
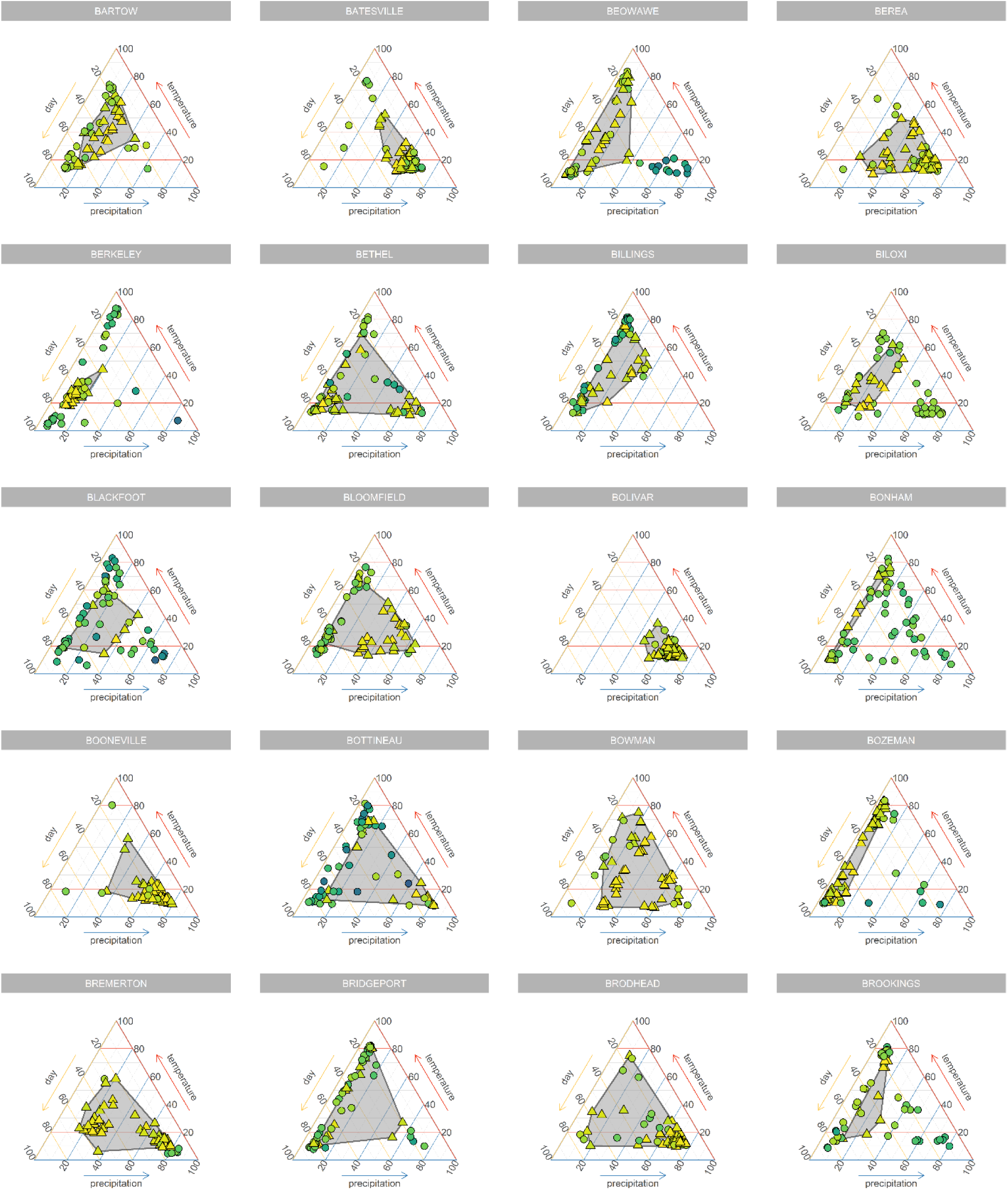
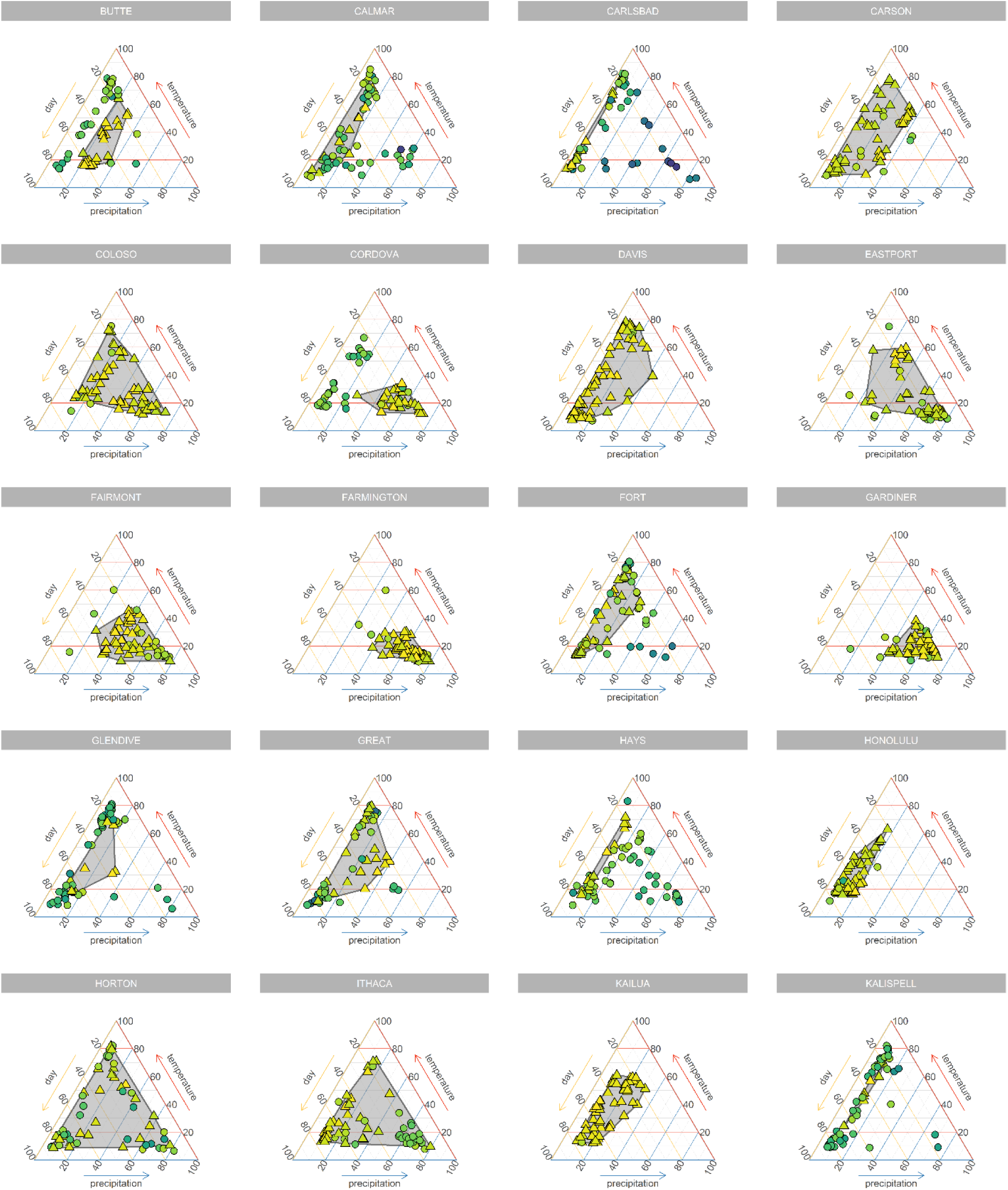
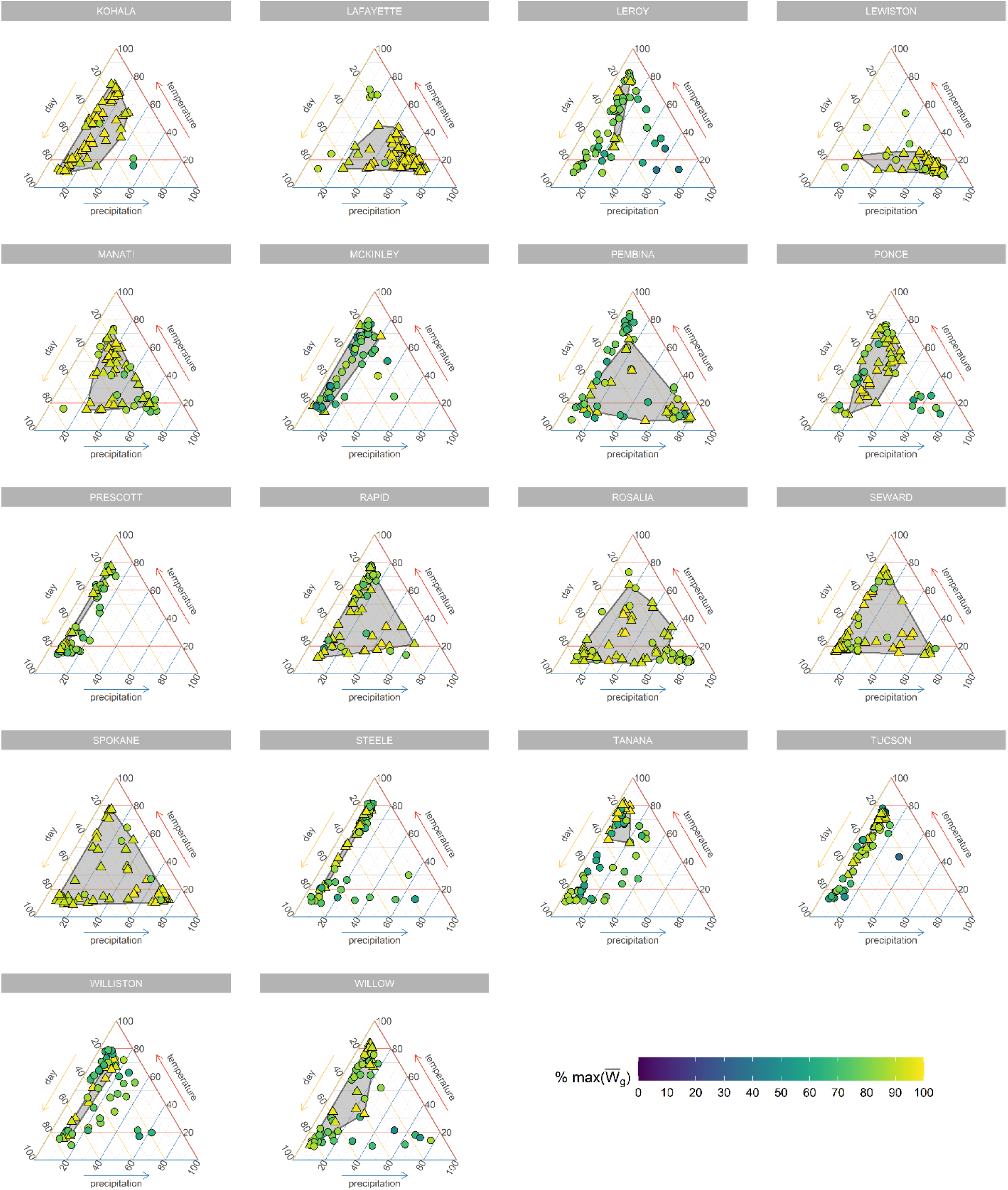
Evolved strategies for 60 simulations in each of 78 locations. Each point represents the mean strategy at the end of one simulation; each strategy is represented as a composition of the “trait effects” in percentages, which represent relative cue use (see “assessing realized relative cue use” in Methods). Point color reflects the mean geometric mean fitness for each simulation scaled by location. Simulations within 10% of the maximum observed geometric mean fitness in each location are represented as triangles and included in a gray convex hull. All other points are represented as circles.

**Figure S6.**
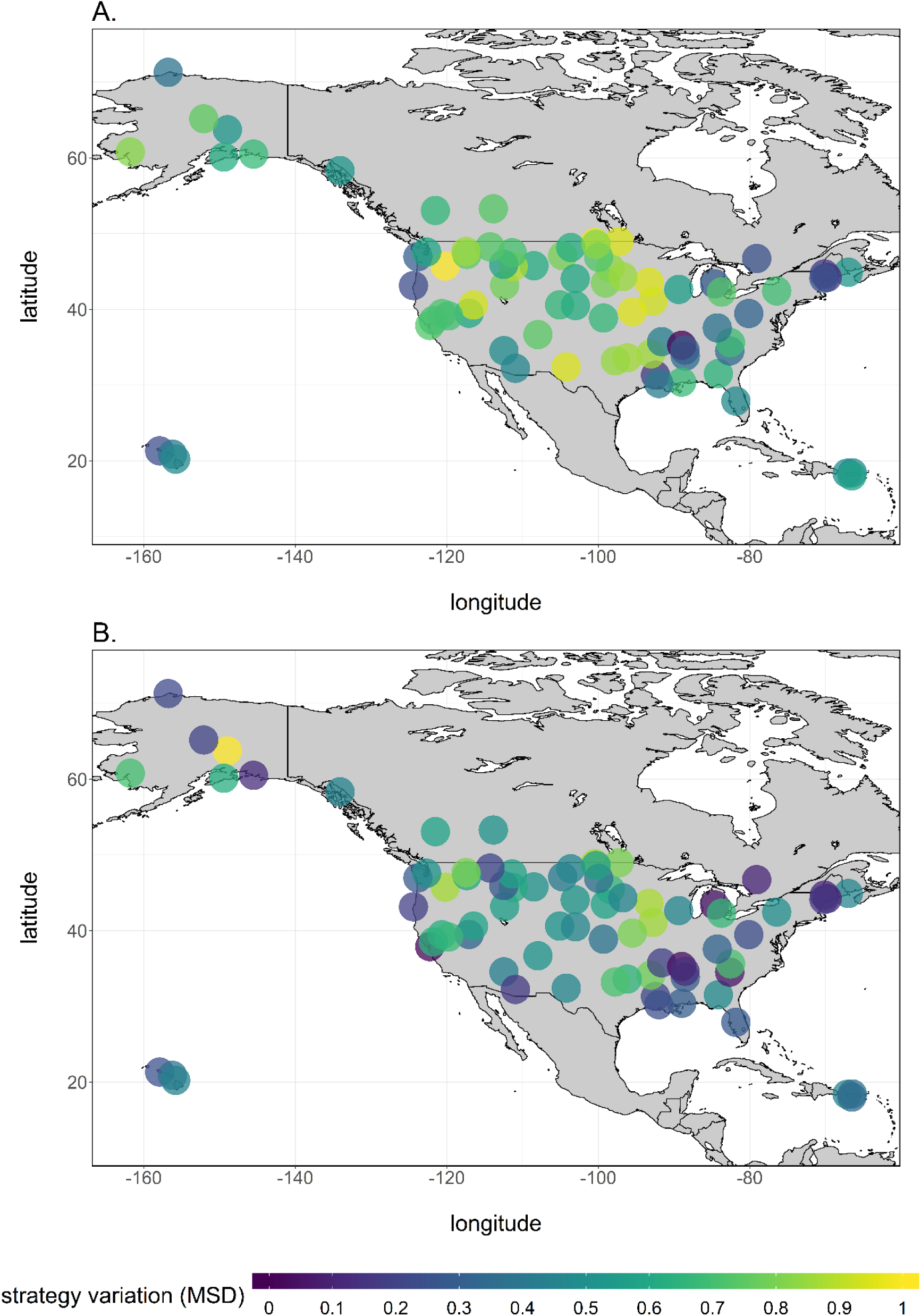
Strategy variation across locations. Strategy variation in each location is measured as the compositional metric standard deviation A) among all genotypes and B) among all genotypes within 10% of the most fit per location.

**Figure S7.**
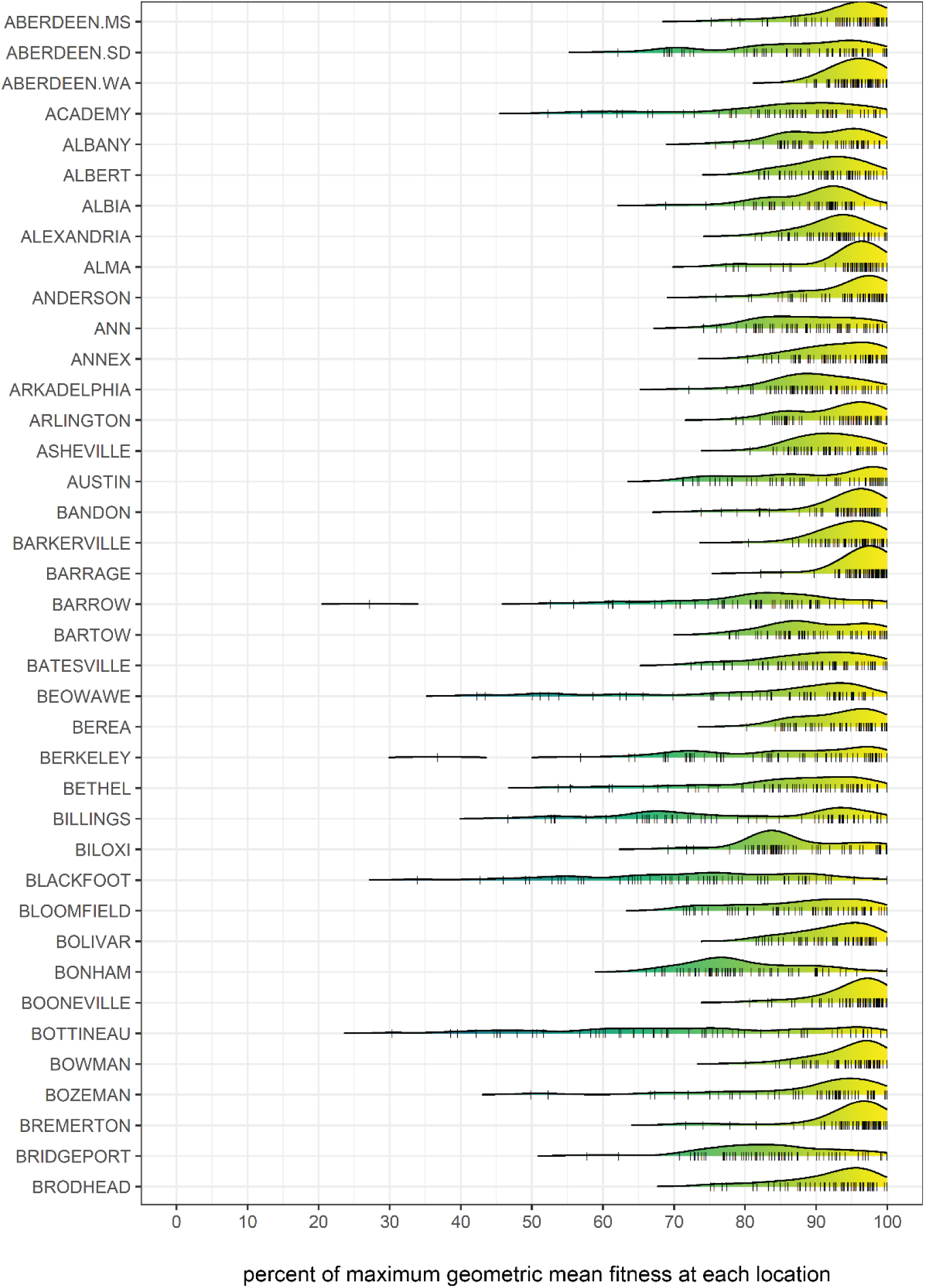
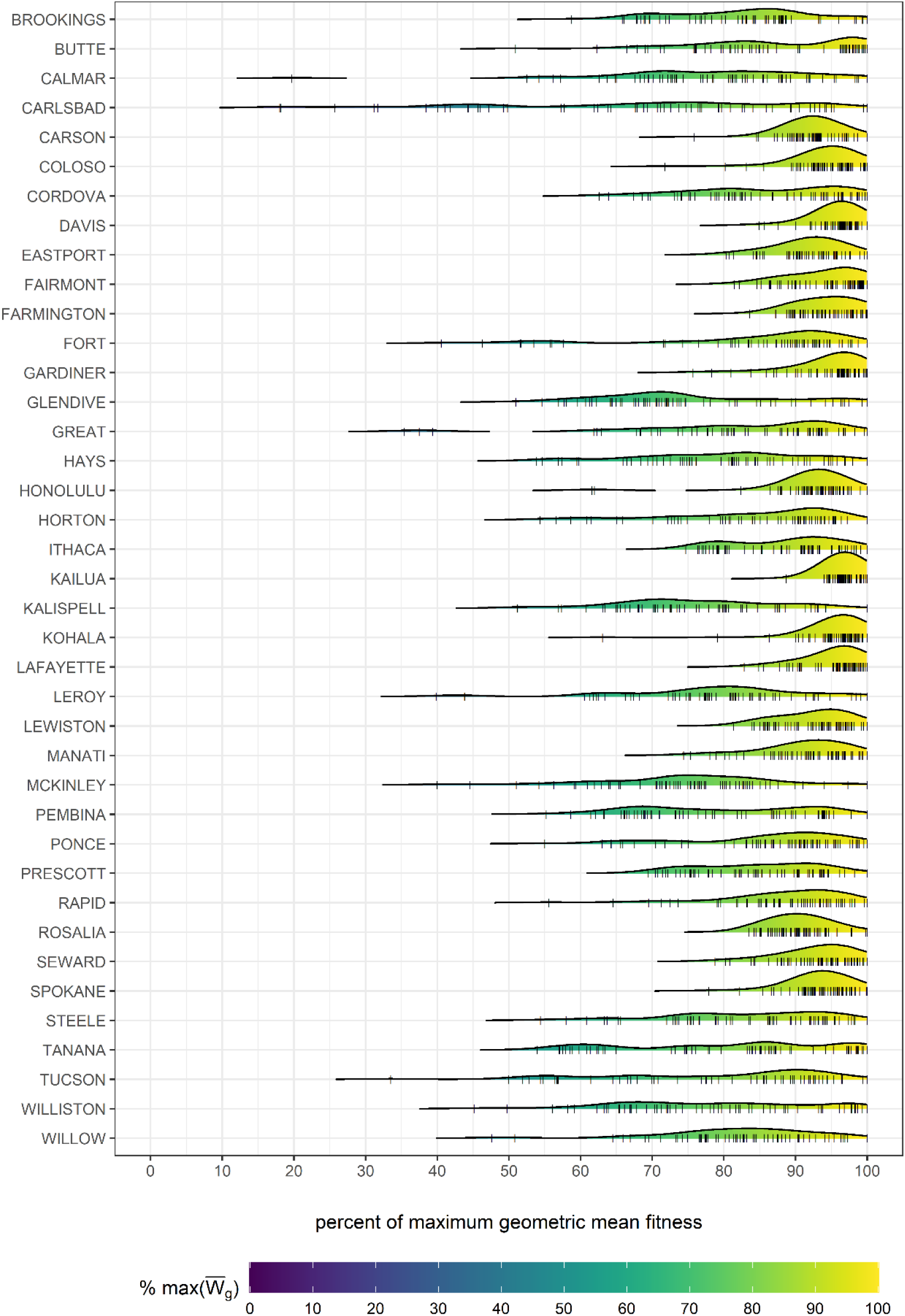
Probability density distributions of geometric mean fitness for 60 simulations in each of 78 locations, rescaled to the most fit simulation in each location. Rug plot marks along the horizontal axis represent individual simulations. Most simulations result in evolved strategies that have geometric mean fitnesses that are within 10% of the most fit simulation in each location.

**Figure S8.**
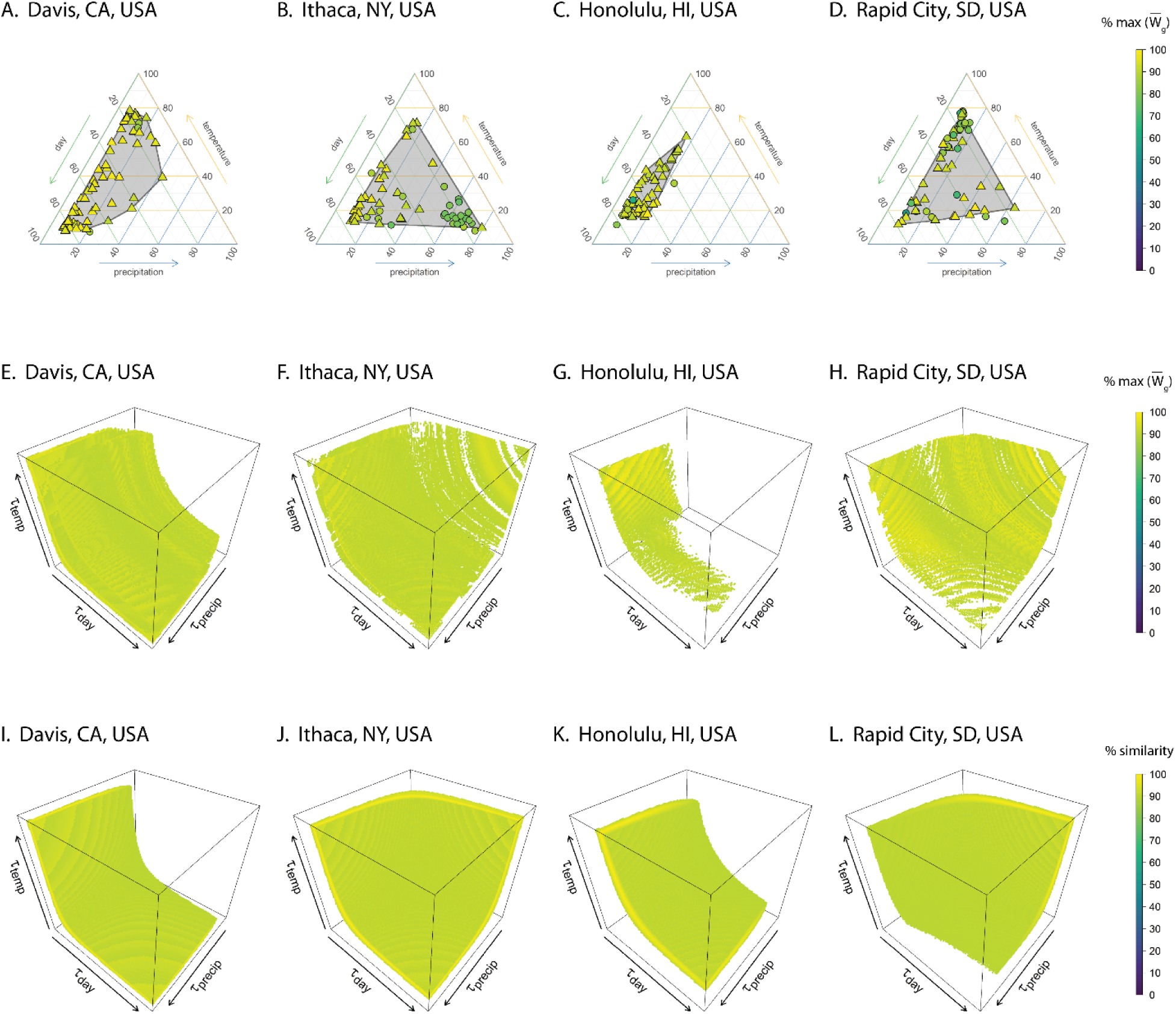
This figure provides an alternative view of the data shown in Fig. 2, where the ternary plots in panels **A-D** are unchanged, panels **E-H** show only the trait combinations within 10% of the most fit combination, and panels **I-L** show only those trait combinations with a response similarity within 10% of the most fit combination.

**Figure S9.**
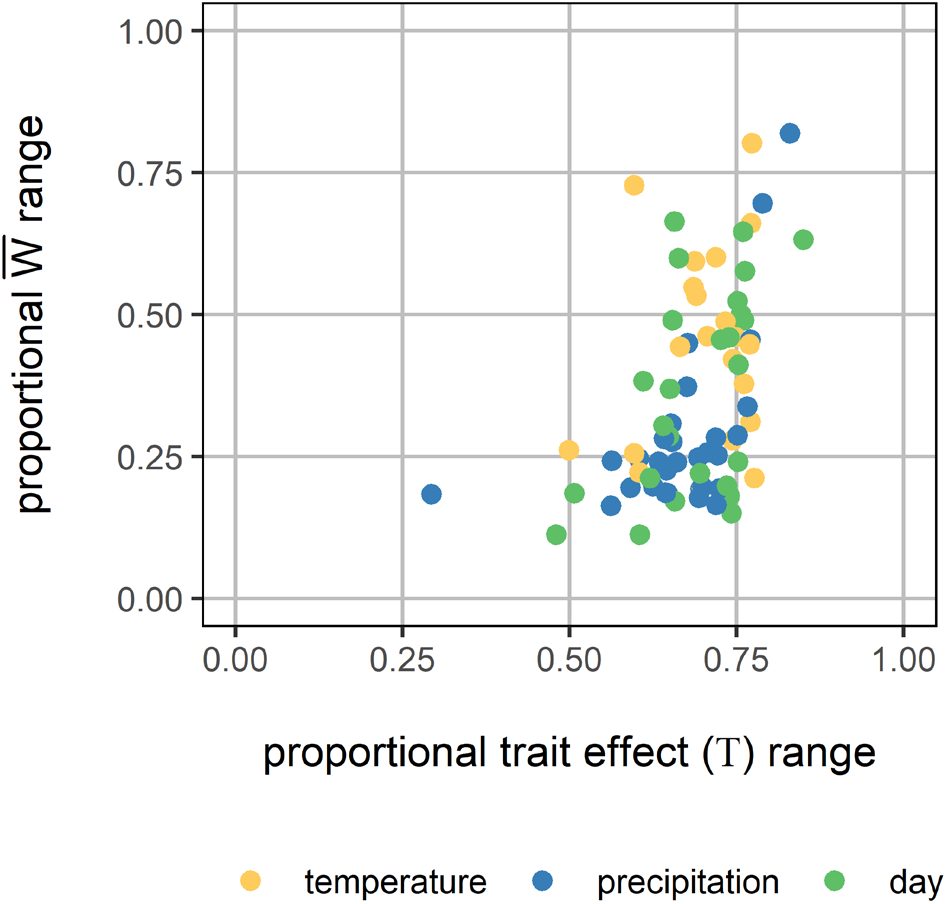
In this figure, each point represents the maximum proportional range of trait effects (Τ) and the maximum proportional range of geometric mean fitness 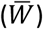 across 60 simulations for each of 78 locations. The maximum proportional range of trait effects is the largest observed difference between the trait effects across simulations within a location, across all three trait dimensions. The color of each point depicts which of the three trait effect dimensions produced the maximum range. Likewise, the maximum proportional geometric mean fitness range is the difference between fitness of the most-fit evolved strategy and the fitness of the least fit evolved strategy. Most locations show a wide range of trait effects with relatively small effects on mean geometric fitness.

**Figure S10.**
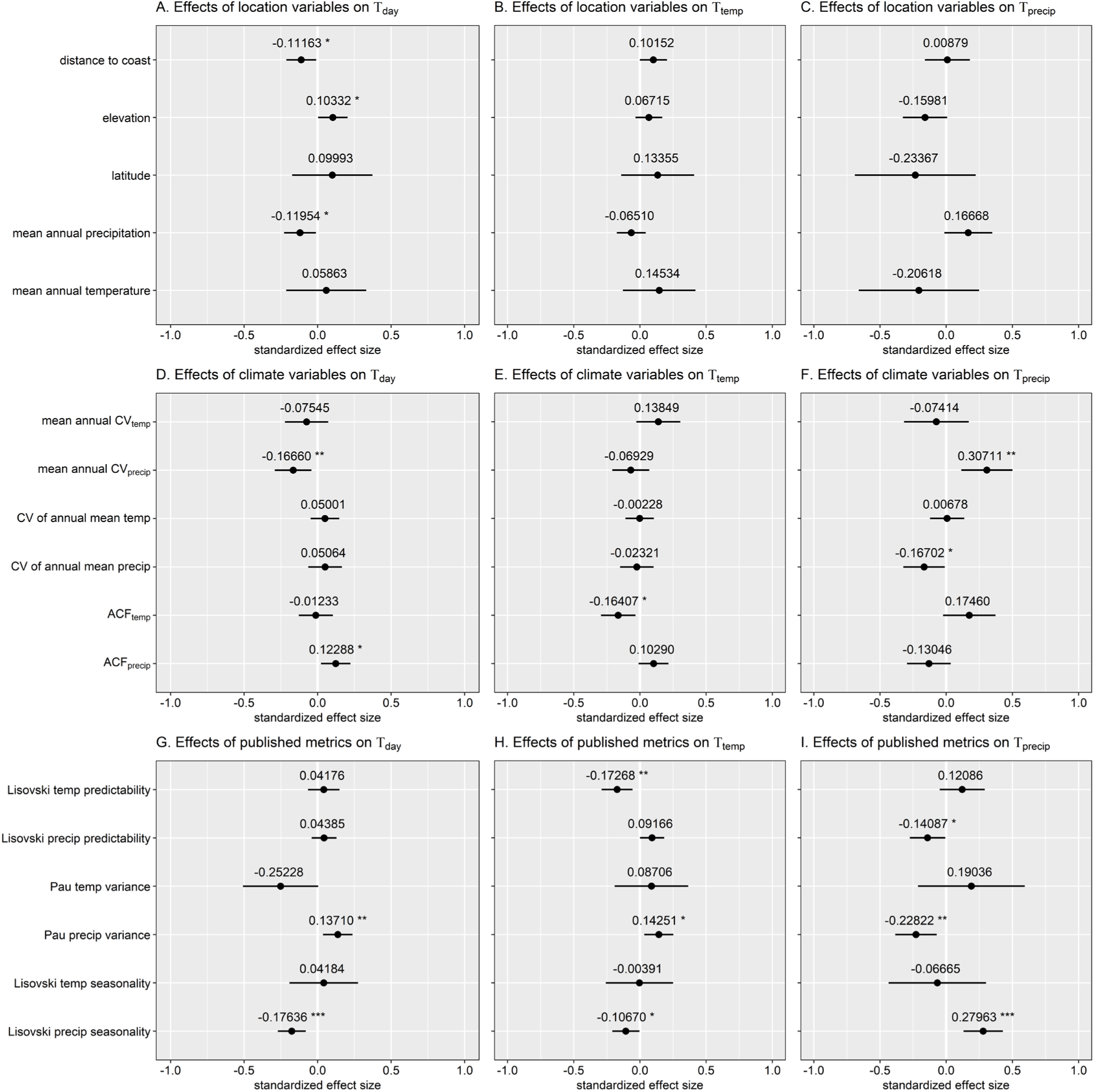
The effects of hypothesized explanatory factors on evolved cue use across all locations under historical conditions. These analyses examine **A-C)** simple location variables, **D-F)** metrics of within- and between-year climate variability, and **G-I)** published metrics of climatic variability, predictability and seasonality. Values indicate standardized fixed effect coefficient estimates, where the raw fixed effect coefficient is divided by the standard deviation of the predictor so that the resulting effect size represents the estimated change in trait effects with a change of one standard deviation in the predictor (see also Fig. S11). Error bars indicate 95% confidence limits.

**Figure S11.**
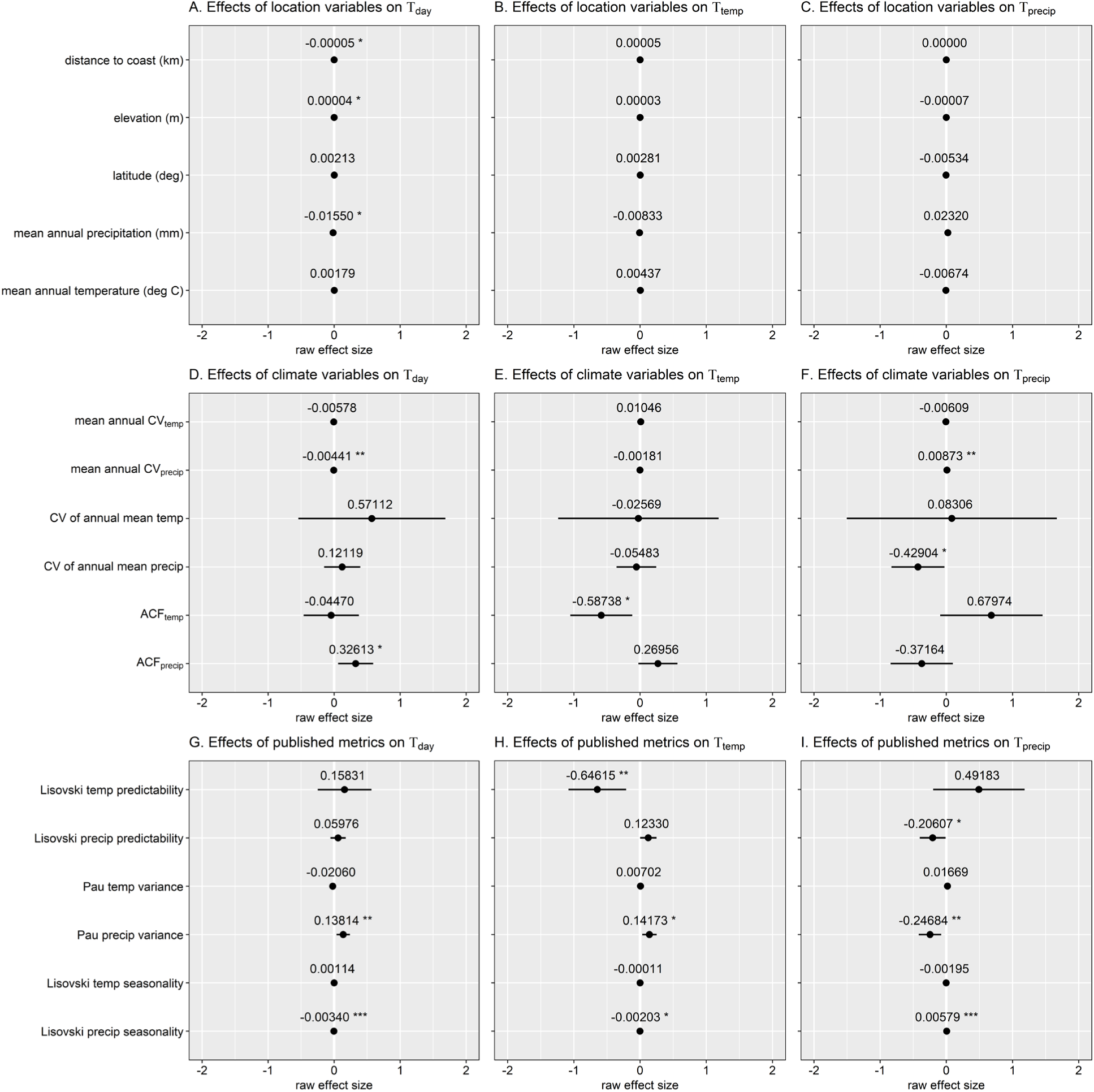
The effects of hypothesized explanatory factors on evolved cue use across all locations under historical conditions. These analyses examine **A-C)** simple location variables, **D-F)** metrics of within- and between-year climate variability, and **G-I)** published metrics of climatic variability, predictability and seasonality. Values indicate raw, unstandardized fixed effect coefficient estimates from linear models (see also Fig. S10). Error bars indicate 95% confidence limits.

**Figure S12.**
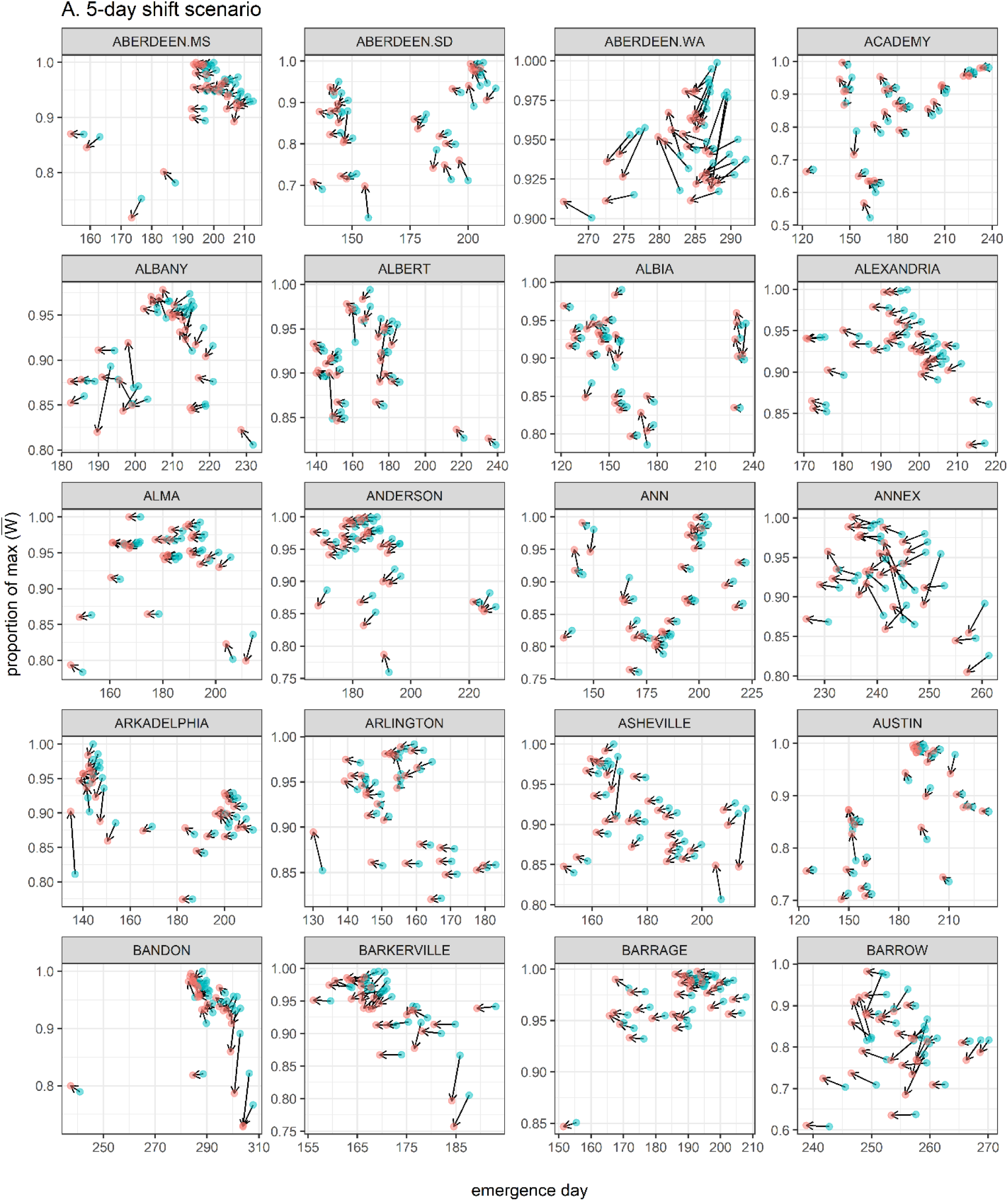
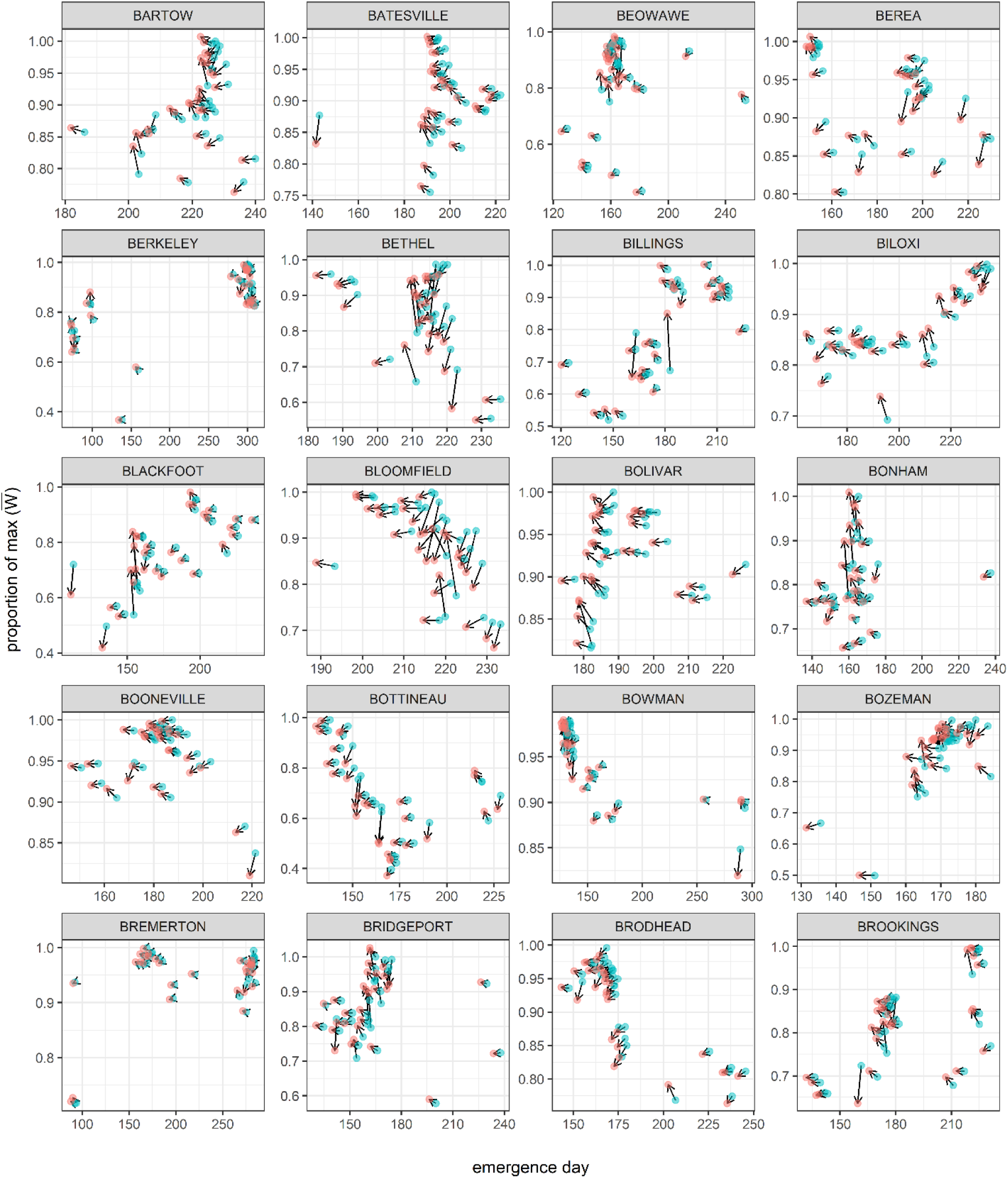
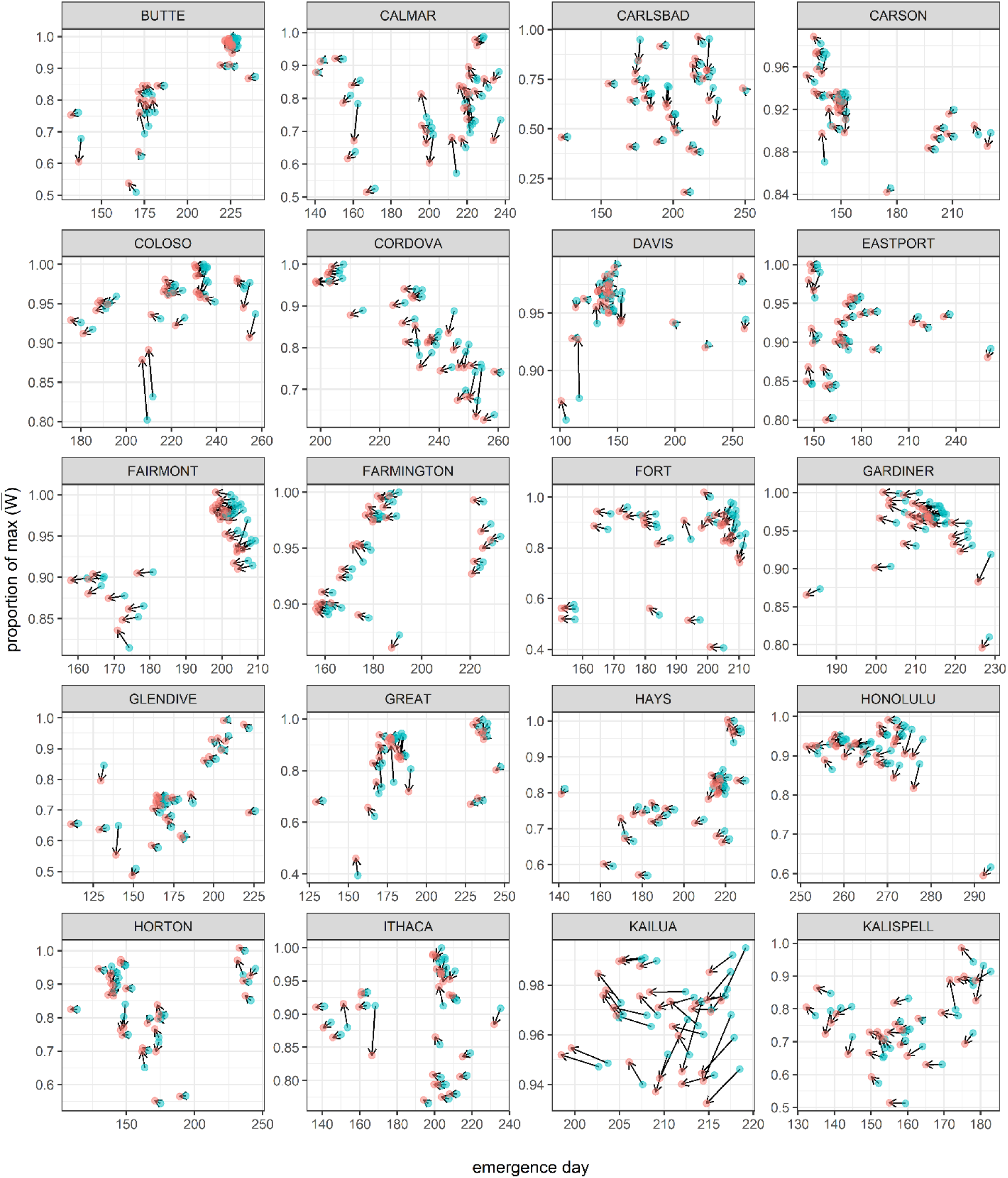
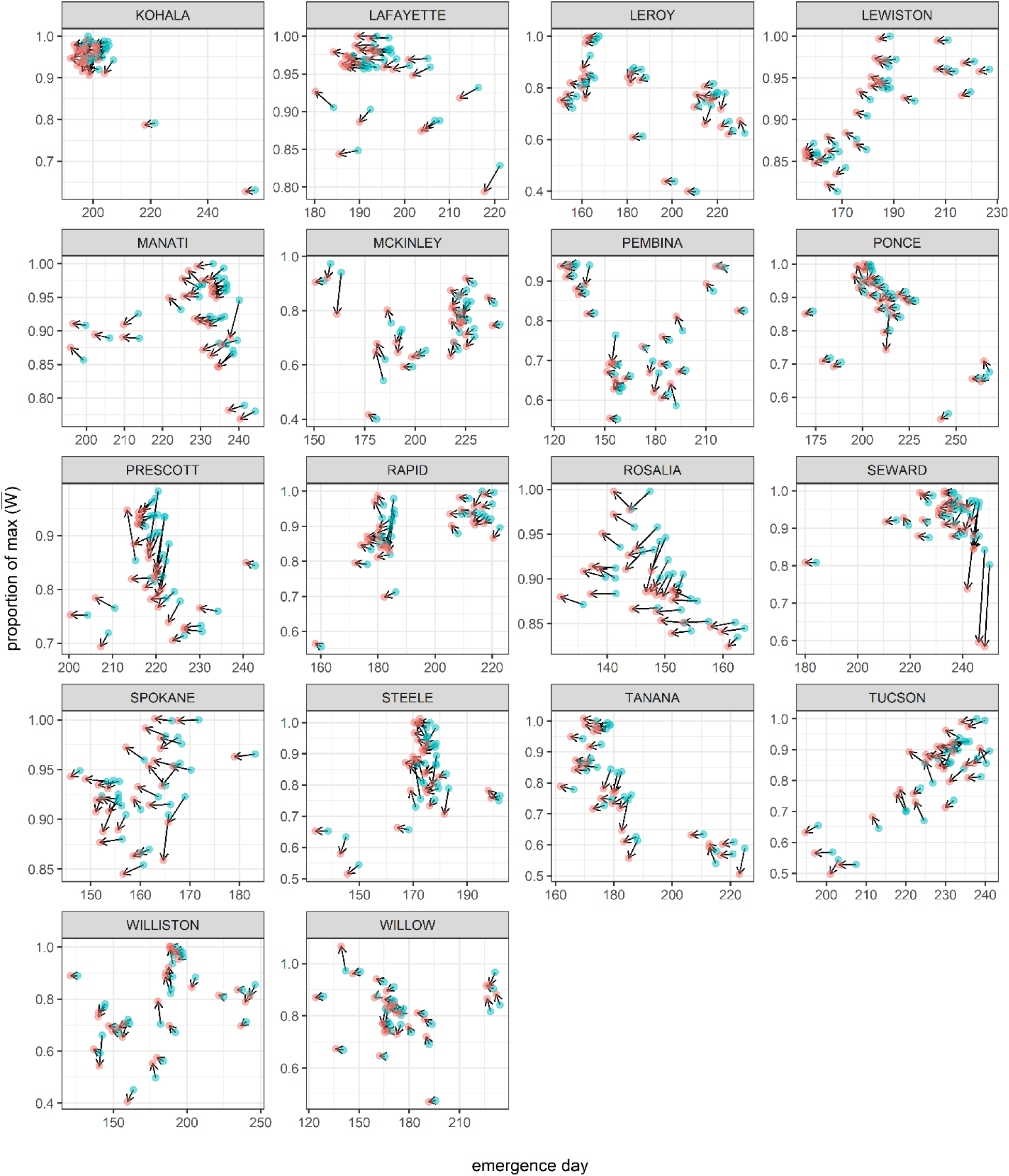
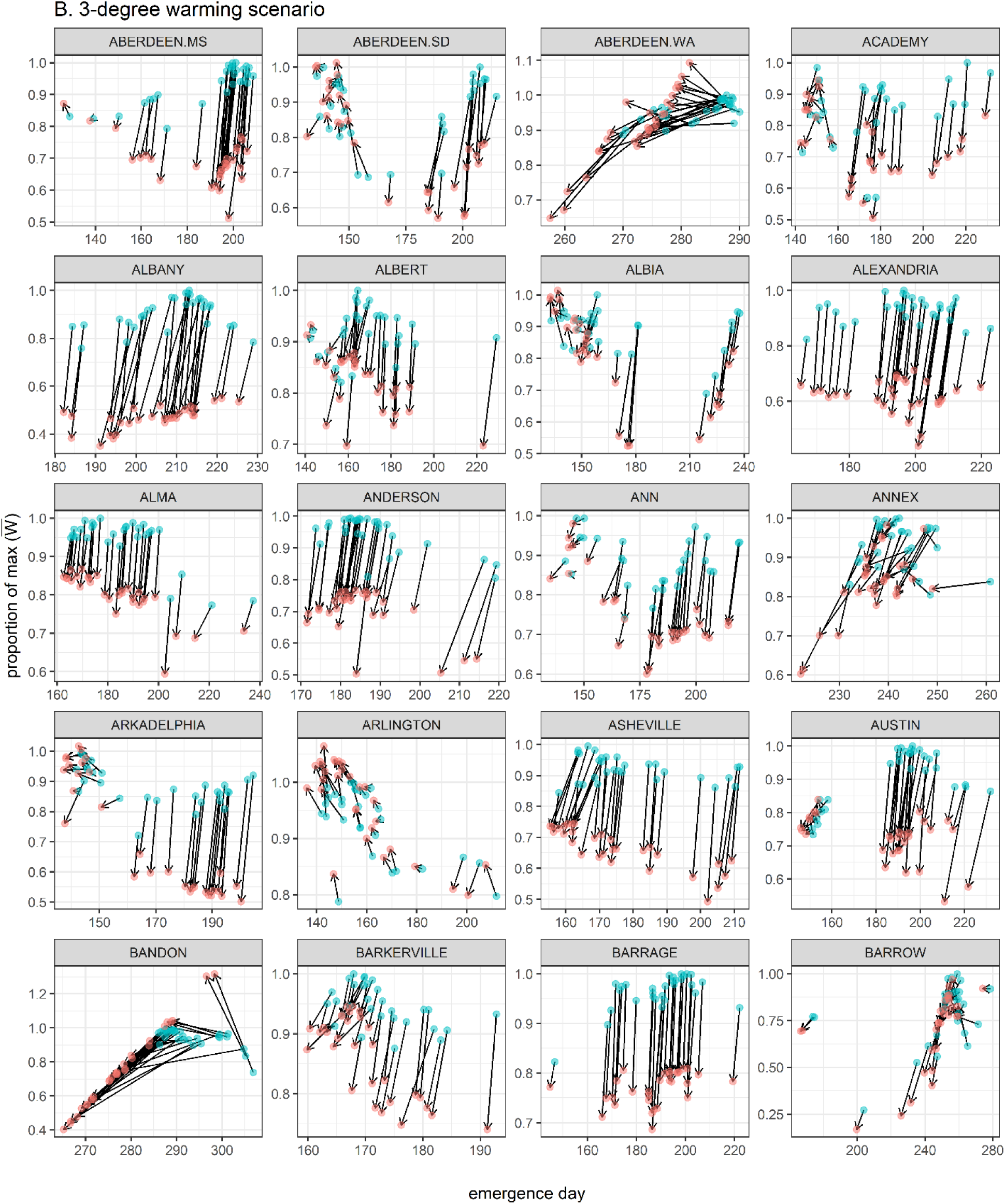
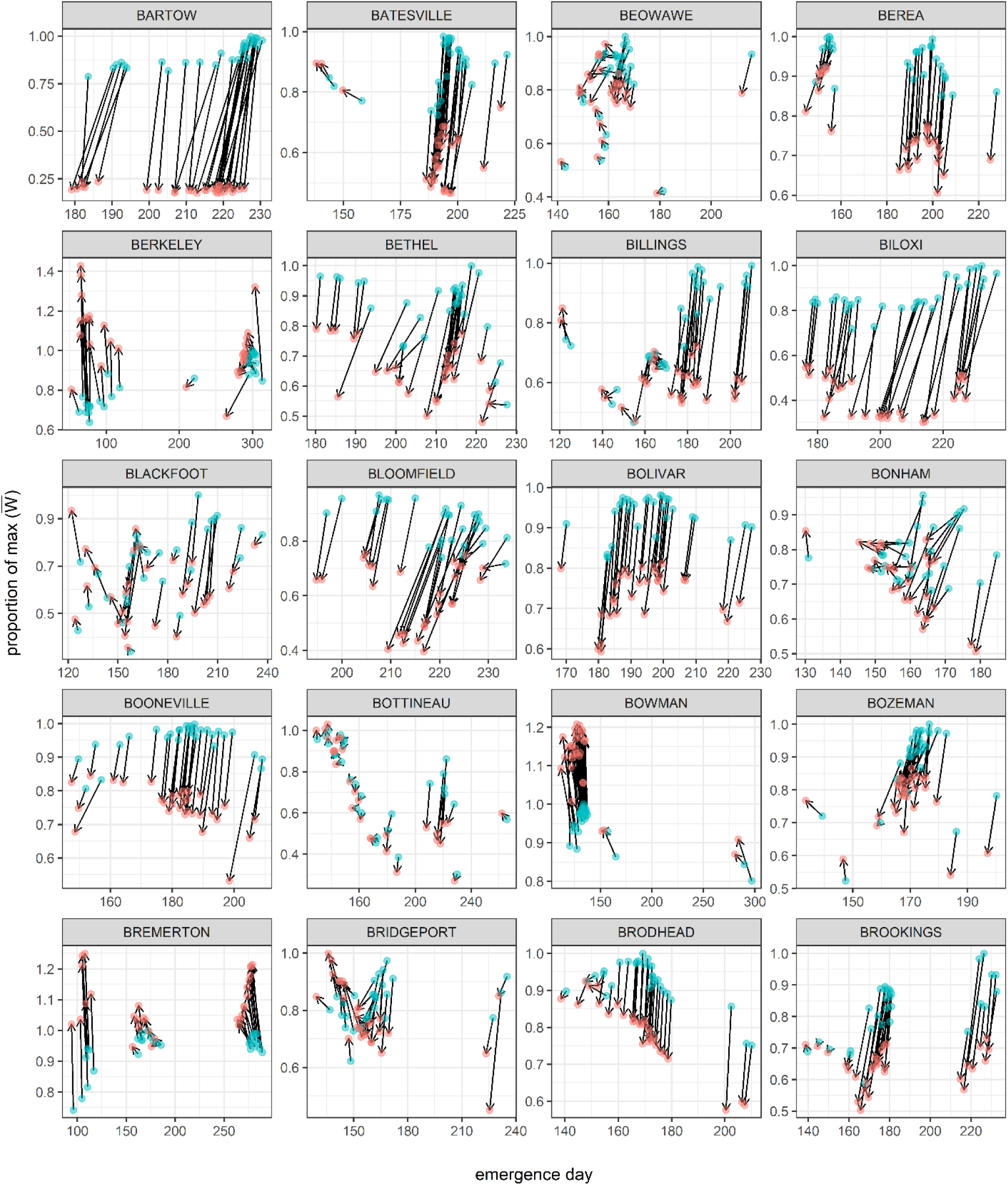
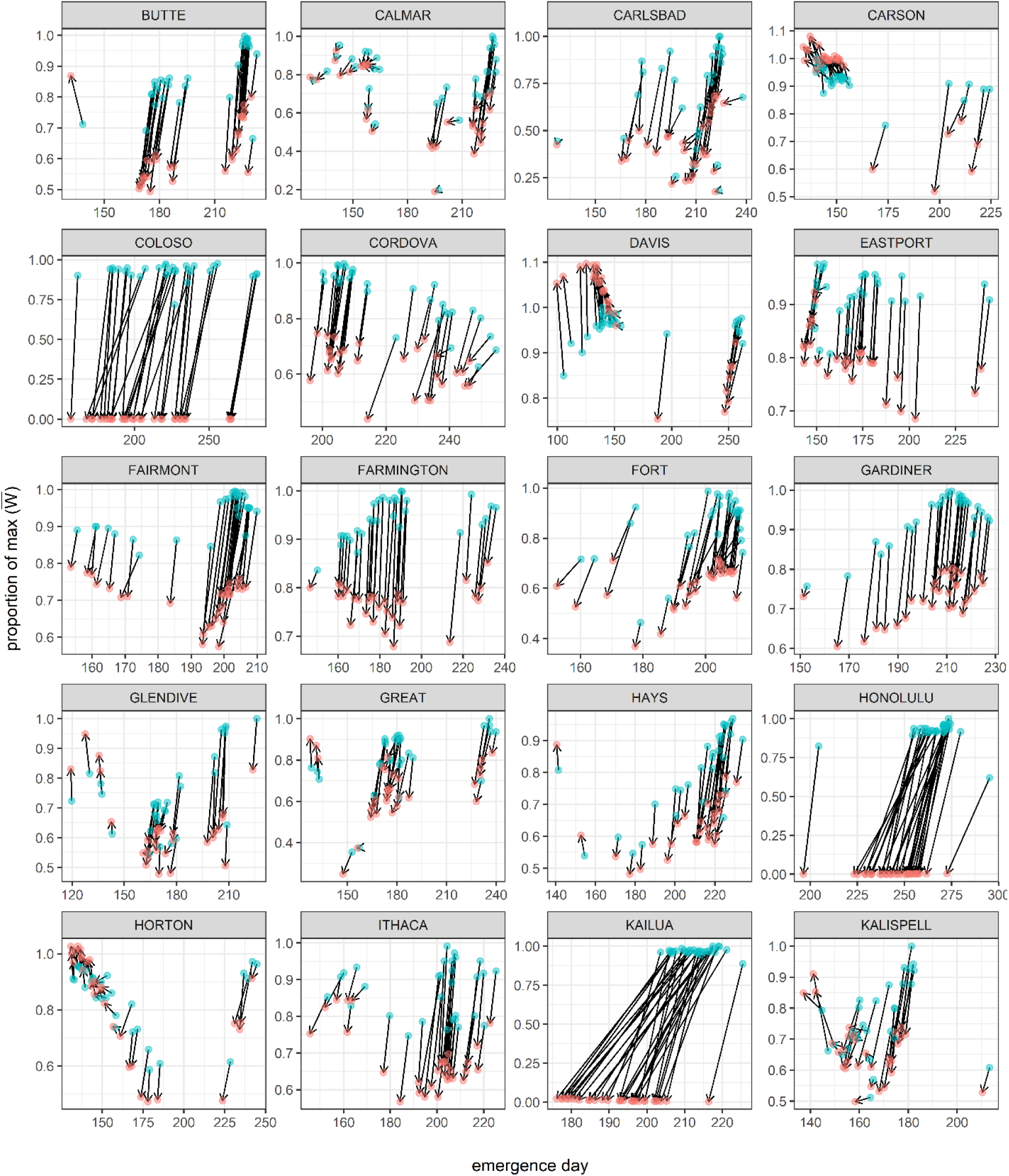
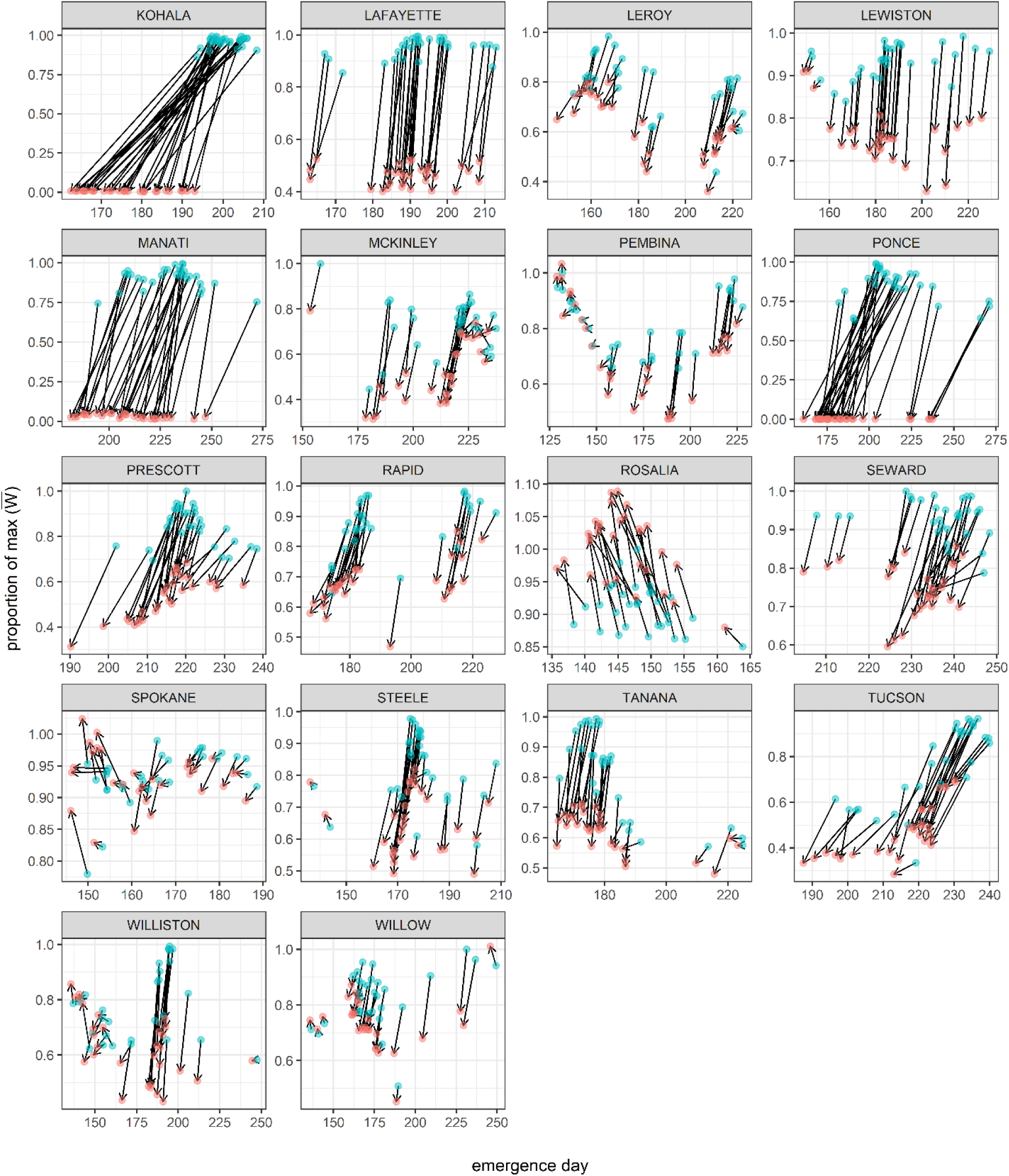
Location specific responses to **A)** a 5-day shift in temperature and precipitation regimes and 3-degree C warming. For each location, each of 30 genotypes is represented by a blue and red circle showing its day of response and proportional fitness under historical and changed conditions, respectively.

**Figure S13.**
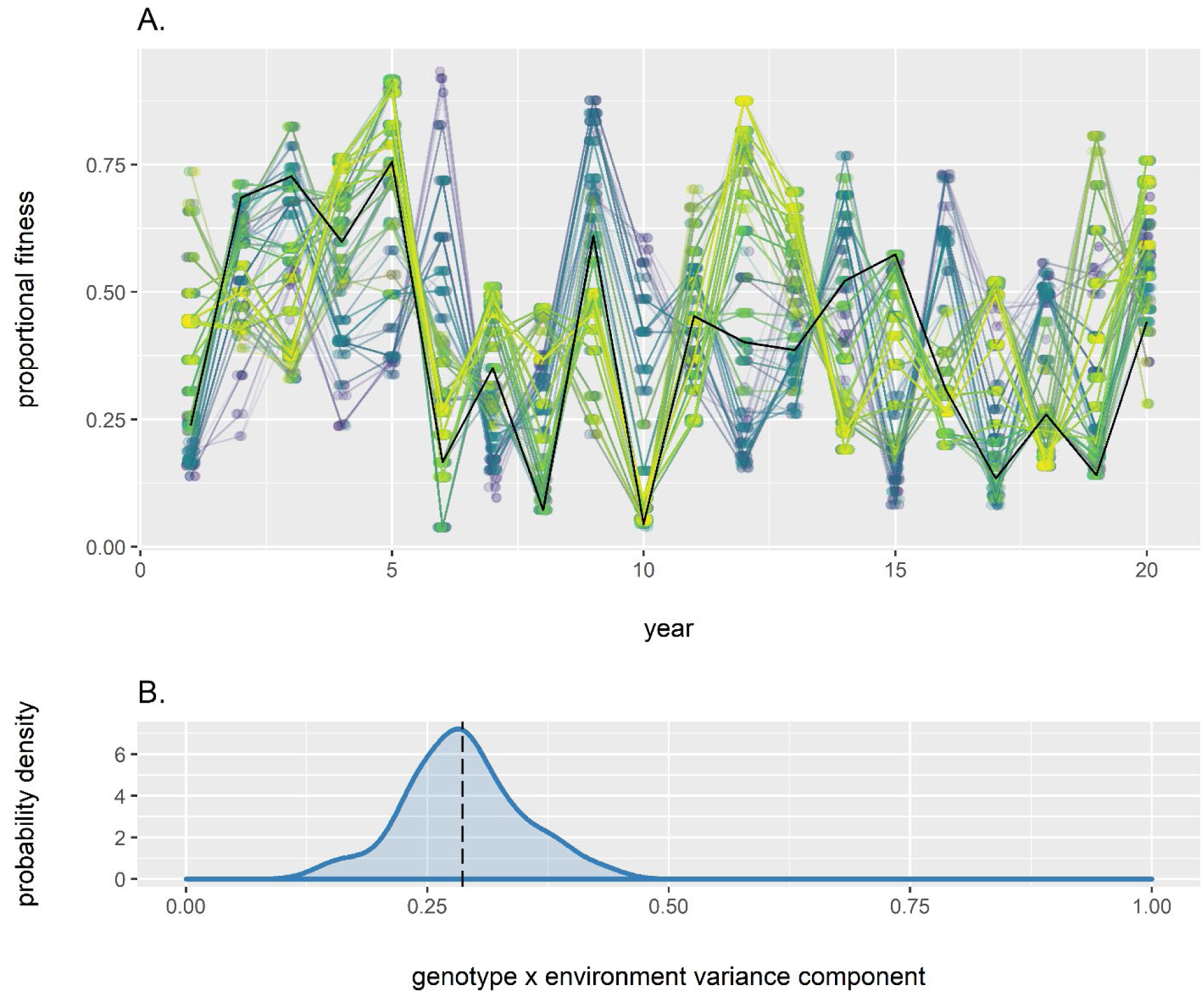
Genotype x environment interactions mean that different genotypes have greater relative fitness advantages in different years; phenological cueing strategies interact with climatic variation to reduce consistent fitness advantages (see Appendix, *“Genotype-by-environment interactions”*). **A)** In this example, each color represents an individual genotype from the final generation of a simulation using climatic data from Ithaca, NY, USA, evaluated in a random subset of 20 years. To facilitate interpretation, a random individual is highlighted as a black line. Fitness has been scaled by the maximum observed fitness across all years for this simulation. **B)** Across all locations, genotype x environment interactions account for 29% of fitness variation on average. The blue probability density function represents the distribution of mean fitness variance components due to genotype x environment interactions in each of 78 locations. The dashed line shows the mean proportion of variation attributable to genotype x environment interactions.

**Figure S14.**
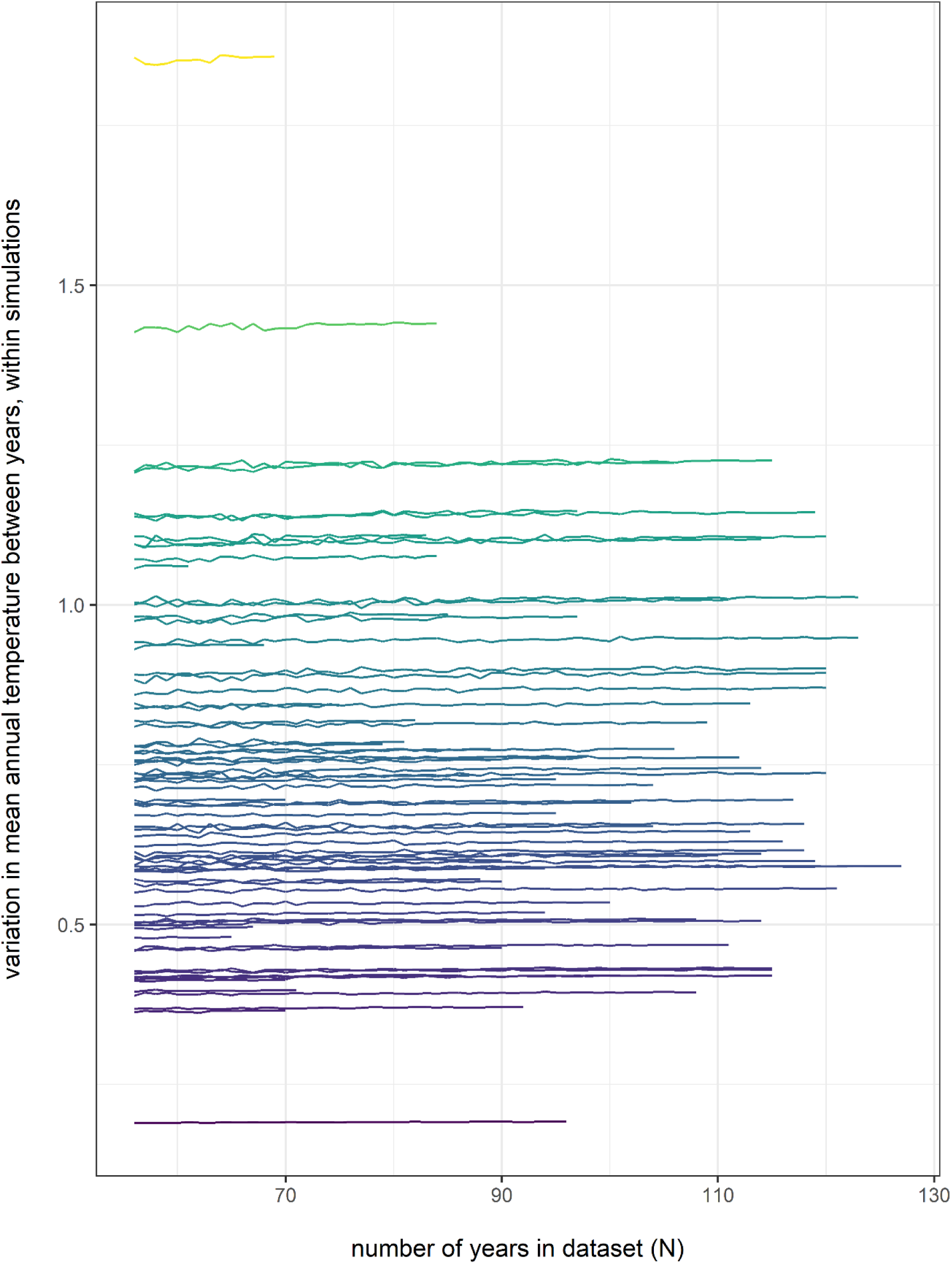
Variation in mean annual temperature between years, within simulations for subsampled climatic datasets of length N, where N ranges from 56 to the full dataset length for each location. N did not have a meaningful effect on variation in mean annual temperature between years, within simulations (Appendix, *“Sensitivity to dataset length”*). Color reflects variation in mean annual temperature between years, within simulations.

**Figure S15.**
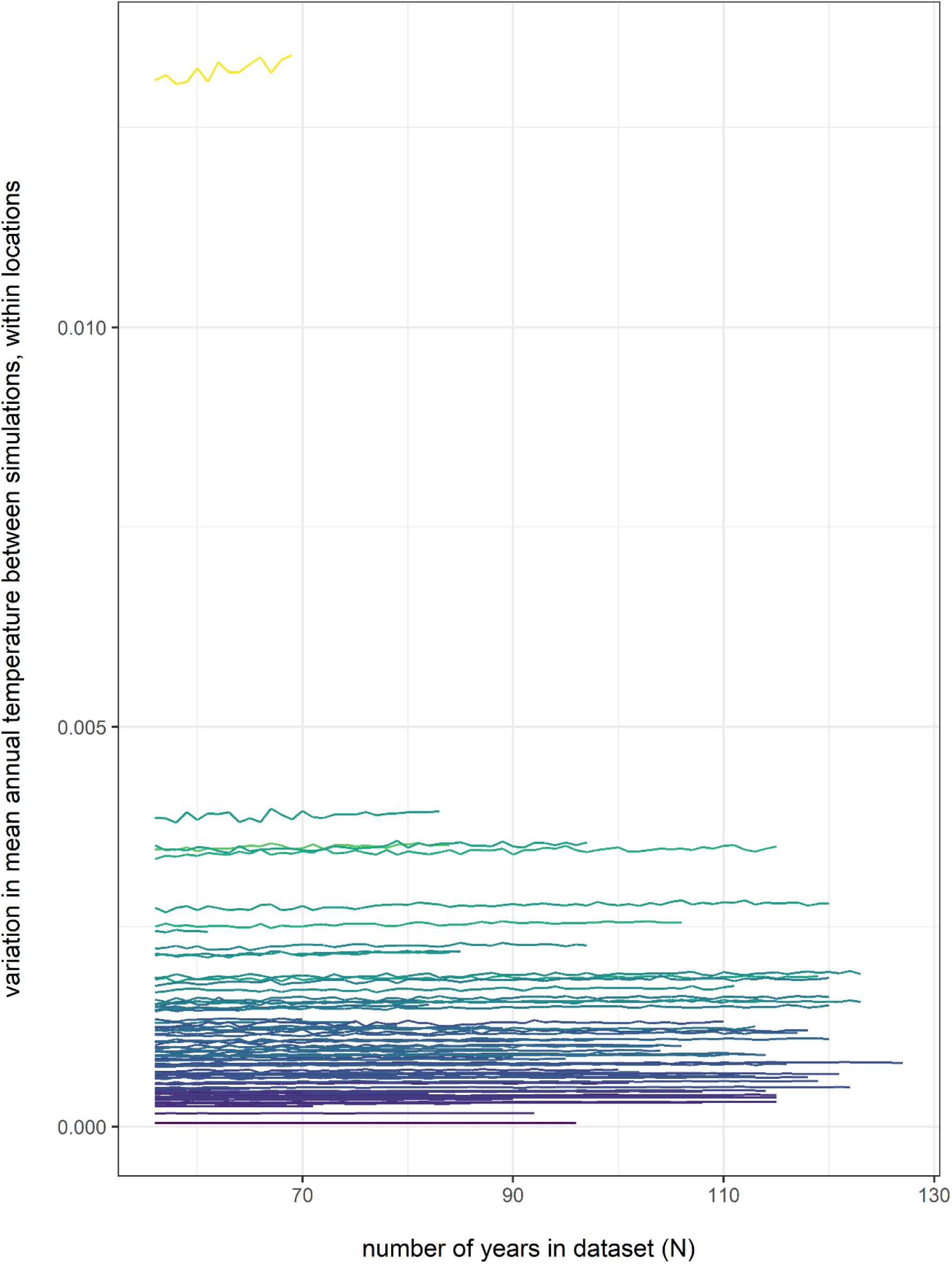
Variation in mean annual temperature between simulations, within locations for subsampled climatic datasets of length N, where N ranges from 56 to the full dataset length for each location. N did not have a meaningful effect on variation in mean annual temperature between simulations, within locations (Appendix, *“Sensitivity to dataset length”*). Color reflects variation in mean annual temperature between years, within simulations.

**Figure S16.**
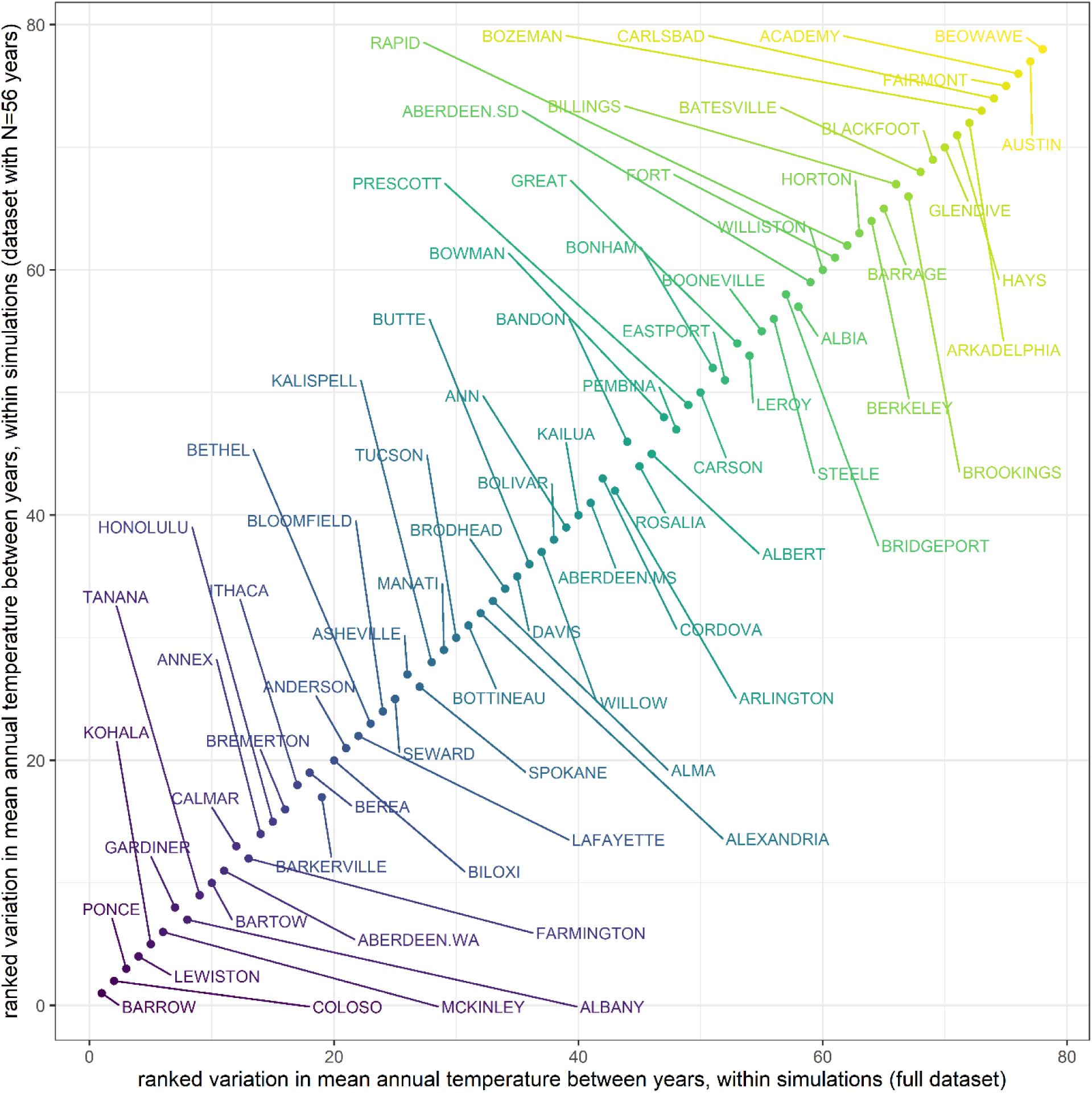
Subsampling from the full dataset length to N=56 years did not meaningfully change the rank order (ranked by variation in mean annual temperature between years) of locations within simulations (Appendix, *“Sensitivity to dataset length”*). Color reflects the rank order of variation in mean annual temperature between years, within simulations for the full dataset.

**Figure S17.**
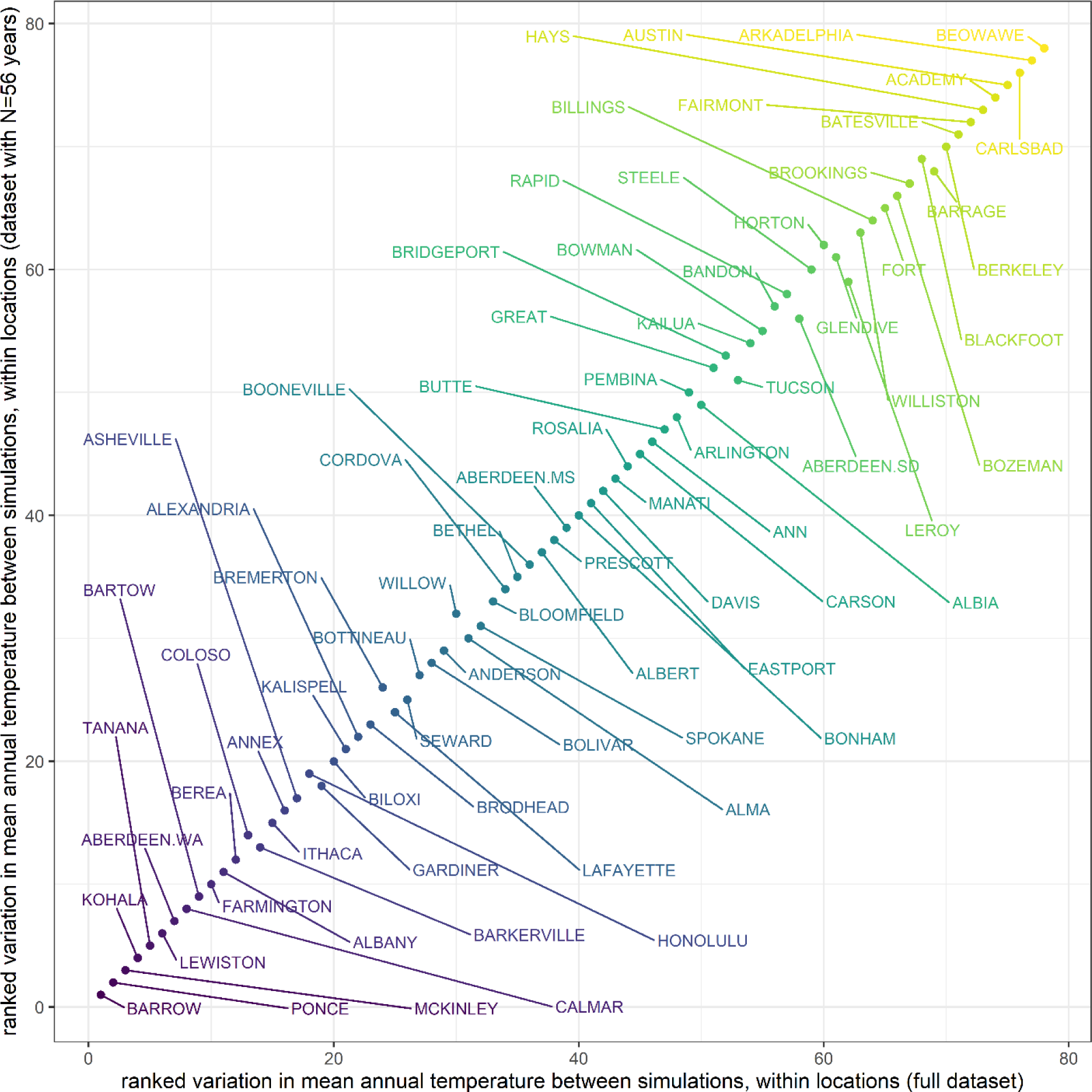
Subsampling from the full dataset length to N=56 years did not meaningfully change the rank order (ranked by variation in mean annual temperature between years) of locations within locations (Appendix, *“Sensitivity to dataset length”*). Color reflects the rank order of variation in mean annual temperature between simulations, within locations for the full dataset.

## Appendix Supporting Methods and Analyses

### Environmental data

Beginning with the raw climatic data from 82 locations in North America and Hawaii, we removed years with <325 daily temperature and precipitation observations, and subsequently excluded four locations with <50 years of data available. For each of the remaining locations, we calculated the interquartile range (IQR) of temperature as the difference between first quartile (Q_1_) and third quartile (Q_3_) observations. Temperature observations less than (Q_1_-4*IQR) and greater than (Q_3_+4*IQR) were identified as outliers likely resulting from measurement error, and were excluded; such outliers were generally climatically impossible, and represented a small proportion (0.0012%) of the overall dataset. Missing observations in the remaining dataset (less than 1% of observations) were imputed using an expectation-maximization with bootstrapping (EMB) algorithm. Imputation was conducted with the expectation-maximization with bootstrapping (EMB) algorithm implemented in the **Amelia II**^1^ package, with priors informed by long-term climatic daily means and standard deviations. Imputation was conducted with the script *“enviro_data_imputation_final.R”*. Imputed values for temperature and precipitation were bounded by the observed minimum and maximum values of each location, and informed by priors based on the means and standard deviations of each location.

For each location, we calculated the minimum distance to a marine coastline using a high resolution coastal shapefile^2^ with the **rdgal** package^3^. Coastal distance was calculated with the script *“distance-to-coast.R”*.

Locations are listed in Supplementary Table 1 along with their ID for use in R data files.

### Sensitivity analyses

We explored several alternative model structures, cues, and parameter values to assess the robustness of our key results. We ran the simulation with both linear and quadratic terms for each of our cues to assess the effects of model structure. We ran the simulation using (a) daily precipitation, daily temperature, and day-of-year, and (b) cumulative temperature, cumulative precipitation, and cumulative photoperiod to assess alternative cues. We ran the simulation on a subset of locations using a product of symmetrical Gaussian distributions, and a product of skew Gaussian distributions with other skew parameter values to test if our simulation was sensitive to the functional form of the fitness surface. We ran the simulation on a subset of locations using larger and smaller population sizes, longer and shorter simulation durations, larger and smaller mutation distances, reduced mutation rates, fitness windows of 5, 10, and 20 days, and higher and lower moisture retention coefficients to test if our simulation was sensitive to these parameter values. We calculated the geometric mean fitness across a broad range of trait values for each location using (a) base parameter values, (b) two alternate parameterizations for each location, each having randomly assigned fitness windows (5, 10, or 20 days), and randomly and independently assigned optimal temperature and moisture quantiles for the fitness function to assess the robustness of our finding that a broad range of genotypes can yield similar fitnesses.

Our general findings were qualitatively robust across model variants that used different cues (daily precipitation, daily temperature, photoperiod, quadratic measures of cues), fitness functions, and fitness windows. Variants with alternative or additional cues also yielded qualitatively similar spatial variation in cue use (e.g, Fig. S1), but each additional cue has a multiplicative effect on potential trait space, and alternative models lacked a strong *a priori* basis. Different fitness window lengths produced qualitatively similar patterns, with longer windows creating more consistent fitness patterns between years (e.g., responding a day earlier or later had a smaller proportional impact on the sum of daily fitnesses). Changing mutation rates, distances, simulation duration, and population size influenced the total amount of trait change that occurred within each simulation (and in the case of duration and population size, dramatically changed computational runtime), but produced qualitatively similarly patterns. Changes to fitness parameters, including optimal temperature and moisture quantiles, did not change the shape of the high-fitness regions in Fig. 4.

We also conducted a full simulation using a 50-day fitness accumulation window in all locations in order to represent a longer-lived organism or an organism with sensitivity to climatic conditions over a broader period of its annual life cycle. The results of this simulation were consistent with our general findings using a 10-day window and with previous sensitivity analyses – this simulation also showed spatial autocorrelation in mean strategies across locations, with substantial variation in cueing strategies within and between locations. This simulation showed two predictable, quantitative differences compared to simulations using a 10-day window. First, evolved strategies under the broader window showed a greater reliance on day-of-year cues. We believe this occurs because the broader window reduces fitness differences between response days and makes coarser seasonal patterns more reliable. Second, when comparing populations within locations, we still see considerable genotypic variation for many locations, but with almost no fitness differences. This further supports our interpretation that the variation we observed between simulations and within locations reflects effective alternate strategies. Thus, these results provide even stronger support for our key findings than the results we present in the main text, due to the relative simplicity of the seasonal fitness landscape under a substantially longer fitness accumulation window.

Thorough exploration and examination of trait-space proved computationally prohibitive, but for each location we evaluated the geometric mean fitness of 5,000,000 genotypes generated from a Latin hypercube implemented in lhs^4^, constrained to the same “reasonable range” of traits we used for the population initialization in the simulations. During this evaluation, we examined how model results changed when we varied lag (the number of days between E≥1 and the organism emerging) between 1 and 5, and the temperature and moisture that maximized fitness between 5^th^ and 95^th^ percentile of observed cue values for each location. Results for these sensitivity analysis were qualitatively similar, but trait values and cue use changed when organisms are rewarded for emerging in low temperatures and moisture. We carried out test simulations in which organisms had different windows (1 and 20 days) after emergence to collect fitness. These simulations showed qualitatively similar results, although simulations with shorter windows were more influenced by extreme weather events, while simulations with longer windows provided smaller benefits for emerging on exactly the optimal day.

### Sensitivity to dataset length

The historical climate data we obtained for our 78 locations varied in length. To evaluate the effect of dataset length on the climatic variation experienced in our simulations, we conducted an additional analysis by repeatedly subsampling each historical dataset without replacement to generate hypothetical data sets with N years of data, where N varies from 56 (the shortest dataset we used) to the full length of the dataset for each location. From these, we replicated the process our simulations used to generate climates for populations: each of the newly generated data sets were sampled with replacement to create 1000-year sequences of climate data, and this process was repeated 60 times for each location. Using mean annual temperature as a representative climate variable, we considered the effects of dataset length on climatic variation at two levels: a) between years, within a simulation, and b) between simulations, within locations. We then plotted both measures of variation against N, where each line represents a different location (Fig.S14, S15). Dataset length had a negligible effect on both measures of climatic variation. We quantified this effect by fitting separate linear regressions for each of these two metrics of climatic variation using dataset length (N) as a fixed factor and location a random factor. The effect size associated with dataset length explained a negligible proportion of the variation a) between years, within a simulation (ΔR^2^ = 0.00004) and b) between simulations, within locations (ΔR^2^ = 0.00003). We also compared the rank order of both variation metrics for all locations using our full dataset versus subsamples of N=56 years, and found that most rankings were unaffected by dataset length (Fig. S16, S17). These results support our assumption that variation in historical dataset length had very little effect on the climatic variation experienced by our simulated populations, and is unlikely to affect the findings of our study.

### Analysis of explanatory factors

We conducted analyses to examine correlations between evolved strategies and three sets of potential explanatory variables at each location. The first set of analyses considered five location variables that provide a broad biogeographic description of each location: distance to coast, elevation, latitude, mean annual precipitation and mean annual temperature. The second set of analyses focused on six variables that quantify climatic variance and predictability: the mean annual coefficient of variation for daily maximum temperature, the mean annual coefficient of variation for daily precipitation total, the coefficient of variation for annual mean daily maximum temperatures, the coefficient of variation for annual mean daily precipitation totals, the lag=1 autocorrelation coefficient for daily maximum temperature, and the lag=1 autocorrelation coefficient for daily precipitation totals. The first two of these variables provide metrics of intra-annual climatic variation, the second two provide metrics of inter-annual climatic variation, and the last two provide metrics of short-term predictability. The third set of analyses used six published metrics of climatic predictability, variability and seasonality (Pau et al. 2011; Lisovski et al. 2017): Lisovski et al.’s predictability and seasonal amplitude (hereafter *seasonality*) metrics for temperature and precipitation, and Pau et al.’s variance metrics for temperature and precipitation. Lisovski’s metric of predictability quantifies the ability of a model parameterized with a moving window of 4 years to predict climatic conditions in the following year across the time series. Pau’s metric of temperature variance is the standard deviation of mean monthly temperatures, and Pau’s metric of precipitation variance is the coefficient of variation for mean monthly precipitation (Pau et al. 2011). Lisovski’s metric of seasonality (i.e., seasonal amplitude) was the difference between the upper and lower 2.5^th^ quantile of the annual distribution (Lisovski et al. 2017). Because the Pau et al. and Lisovski et al. metrics of seasonality were highly correlated, we did not also include the Pau et al. metrics of seasonality.

These analyses used a dataset composed of the mean strategies that evolved in each simulation conducted in each location. For each analysis set, we used linear mixed models including all *a priori* explanatory variables as fixed factors for each trait effect dimension, with an additional random factor to allow intercepts to vary by location. For each analysis set, we ran the linear mixed model separately for each of the three cues. We find qualitatively identical results using logit-transformed trait effects, and primarily present analyses of untransformed data here so that the effect sizes are reported in interpretable units. Standardized effect sizes (B) are represented by the fixed effect slope coefficient divided by the standard deviation of each explanatory variable to allow for comparisons between explanatory factors (Fig. S9); these effects sizes reflect the change in trait effects across one standard deviation of the explanatory variable (Bring 1994). We also present non-standardized effect sizes for completeness (Fig. S10).

Several location-based explanatory factors were associated with variation in phenological cueing strategies under our specific model (Figs. S8). For example, the evolved use of day cues (measured as the mean trait effects of the final generation) declined with distance from the coast (B=-0.112), and increased with elevation (B=0.103). The use of day cues may have been favored in more moderated coastal climates if these climates tend to show more predictable seasonal variation where climatically plastic cues are less relevant for identifying successful windows of opportunity. Conversely, the greater use of day cues may have evolved at high elevation locations if more variable day-to-day climatic conditions meant that climatic cues provided less predictive information than a climatically invariant day cue. In comparison, these two explanatory factors suggest that the evolution of phenological cues may reflect a key distinction between informative climatic variation and noise. In addition, locations with higher mean annual precipitation tended to rely on day cues less (B=-0.120), and generally showed a greater emphasis on precipitation cues (B=0.167). This pattern could result from the absence of informative cueing information provided by precipitation cues in relatively dry locations.

Some metrics of climatic variation also explained the evolved use of phenological cues. Locations with high mean annual CV_precip_ (i.e., high within-year precipitation variability) tended to show a reduced use of day cues (B=-0.166), and an increased use of precipitation cues (B=0.307). In contrast, locations with high CV of mean annual precipitation (i.e., high between-year precipitation variability) tended to show a reduced use of precipitation cues (B=-0.167). These patterns suggest that the use of precipitation cues allows for adaptive plasticity in locations with variable within-year precipitation regimes, but a reliance on precipitation cues is not favored in locations with high between-year variation in precipitation, where precipitation cues may be lacking or uninformative in some years. Unexpectedly, locations with high lag 1 autocorrelation in temperature (ACF_temp_) actually tended to use temperature cues less (B=-0.164), while locations with high ACF_precip_ tended to use precipitation cues less (B=-0.130), and day cues more (B=-0.123). This result is counterintuitive, since it shows a negative correlation between a direct measure of (daily-scale) climatic predictability and evolved cue use. However, a similar pattern emerges from an independent published metric of predictability on a seasonal scale: locations with high Lisovski temperature predictability showed a reduced use of temperature cues (B=-0.173), and locations with high Lisovski precipitation predictability showed a reduced use of precipitation cues (B=-0.141). We speculate that these patterns occur because the use of day cues can compensate to some degree for the use of climatic cues in locations with highly predictable climatic conditions, and because maximizing the fitness outcomes of strategies in locations with one highly predictable climatic condition may depend more on identifying windows of opportunity constrained by the other climatic factor. Pau’s metric of precipitation variance was associated with an increased use of day (B=0.137) and temperature (B=0.143) cues, and a reduced use of precipitation cues (B=-0.228), while Lisovski’s metric of precipitation seasonality was associated with a reduced use of day (B=-0.176) and temperature (B=-0.107) cues, and an increased use of precipitation cues (B=0.280). These patterns are in the opposite direction because the Pau measure of precipitation variance uses the coefficient of variation, and so is negatively correlated with mean annual precipitation (r=-0.54, p<0.0001)), while Lisovski’s metric of precipitation seasonality was highly positively correlated with mean annual precipitation (r=0.94, p<0.0001). The pattern with Lisovski’s measure of precipitation seasonality supports the previous observation that the use of precipitation cues is positively correlated with intra-annual precipitation variation. Overall, while several factors emerged as meaningful predictors of phenological cue use in these analyses, most of the variation in cue use was unexplained even in models that combined all 17 factors for each trait effect (Τ_day_: marginal R^2^=0.102; Τ_temp_: marginal R^2^=0.110; Τ_precip_: marginal R^2^=0.308).

We suggest that the limited success of these explanatory factors for predicting cue use occurs because they are too far removed from the specific selection criterion implicit in the model. For example, metrics that attempt to summarize inter- or intra-annual climatic variation are only distantly correlated with the predictability of cues in the most relevant period of the year when fitness is highest. Likewise, even metrics that specifically target climatic predictability (as opposed to variation) were generally unable to provide a strong explanation for evolved cues because they quantify mean predictability across the year, while selection favors strategies that are able to identify potential windows of opportunity in specific parts of each year. In general, the evolved strategies in our model offers only partial support for conventional expectations based on location characteristics or standard metrics of climatic variability (Sunday et al. 2010; Molina-Montenegro and Naya 2012); instead, they illustrated the complexity of the climatic regimes and seasonal fitness landscapes that affect organisms in this model. Further, the considerable variation in evolved strategies within many locations suggests that simple climatic and environmental factors alone are unlikely to adequately predict the evolution of phenological cueing strategies.

The quantitative results of this analysis are specific to the model parameters we used, and shouldn’t be used to infer the drivers of phenological cue evolution more generally. However, the results of this analysis do suggest several potentially general patterns. Even when the specific relationships are complex or counterintuitive, the observation that some location and climatic variables are predictably associated with reliance on specific phenological cues suggests that these evolved phenological cueing strategies were a predictable outcome of their environmental histories. Moreover, evolution appeared to favor cues that were reliably present from year to year, but showed an intermediate level of variation within year – neither so predictable that they could be replaced by climatically invariant day of year cues, nor so variable that they diluted informative signal with uninformative noise. Thus, the spatial autocorrelation of evolved strategies suggests that past environmental regimes shaped the evolution of phenological cueing strategies in fundamentally predictable ways, even if our ability to characterize the specific environmental factors that favor different cueing strategies is limited.

### Genotype-by-environment interactions

The GxE interactions in our model reflect that the effects of different genotypes (i.e., combinations of three traits, or phenotypic cueing strategies) on phenotype (i.e. day of emergence) or fitness are different in different environments (i.e., years). Broadly, the GxE effect weakens the potential for consistent fitness differences among genotypes (i.e., phenological strategies), promoting the persistence of genotypic diversity (Falconer 1952; Gillespie and Turelli 1989). We quantified the GxE interaction effect by partitioning variation in the annual fitness of a non-evolving population (among individuals, within each simulation, with no mutation or selection) exposed to every year of climate data from that location using a linear regression with fitness as the response variable and individual and year as fixed effects. Because our model is deterministic (i.e., there is no measurement error), the proportion of variation that could not be explained by individual and year factors represents the GxE effect. We calculated the percent fitness variation explained by GxE effects for each simulation as (1 - r^2^) ×100, and report the average percent variation explained by GxE effects for all simulations for all locations.

The genotypes in each simulation of our model show different reaction norms across years, driven by between-year differences in the historical climatic data (Fig. S13A). We estimated that the GxE effect would explain 29% of fitness variation on average across all simulations and locations (Fig. S13B).

### Alternative climate change scenarios

We also tested a “warming + shift” climate change scenario in addition to the two scenarios presented in the main manuscript. This third scenario included a 3-degree increase in daily temperatures, and a shift in the precipitation regime that matched the median phenological advancement in cumulative temperatures. To do this, we calculated the median cumulative temperature value for each location. For each year, we determined the day the cumulative temperature matched or exceeded this value in the historic climate regime and in the shifted temperature scenario. The difference of those two days represented the effective temperature advancement, and we shifted the precipitation regime forward to match this advancement, treating each year as cyclical to accommodate the beginning and end of each year. This produced qualitatively similar responses in the organisms, but this “warming + shift” scenario was complicated by unpredictable shifts in precipitation which varied by location (e.g., warm locations had smaller shifts than cold locations, since the relative change in cumulative temperature from a 3 degree warming was smaller).

### Code guide

The simulation code for this project is designed to work with the following file structure, which is necessary for the code to run. Files are italicized while folders are not. Exact file names are contained in quotes. A ZIP file with the scripts and data files in the appropriate file structure can be found on Dryad at https://doi.org/10.25338/B8TG95.
- Project directory
  - Climate data
    - *“location summary with distance to coast.csv”*
  - data-years
    - *all data files to be used*
    - *“location summary with photoperiod.Rdata”*
  - data-years-lisovski
    - *all data files to be used*
  - fitcurve
    - *“skewgauss.R”*
  - parameters
    - Any parameter files to run. The following are the parameter files used to generate the results presented here:
    - *“cu-ccshift-mut005-parameters.R”*
    - *“cu-cctemp-mut005-parameters.R”*
    - *“cu-cuphotoperiod-mut005-parameters.R”*
    - *“sensitivity-win50-mut005-parameters.R”*
  - results
  - scripts
    - *“master-script.R”*
    - *“rate-setup.R”*
    - *“simulation.R”*
    - *“windows-subs.R”*
  - yearinds

Code is initiated by running *master-script.R*. The “set_wrkdir()” function at the beginning of this file should be updated to point to the home folder you are using, and “prefix” and “suffix” should be updated to match the name of the parameter file to run (by default, set “prefix” to everything before the “.R” of the parameter file name, and set “suffix” to be an empty string). The “sum.merge.only” and “test.only” parameters are to analyze existing runs and test the code using the first two locations, respectively. These should be set to FALSE for a complete run. Outside of the parameter files, these are the only lines of code that may require modification to implement our model.

After reading in parameter (below) and function files, *“master-script.R”* calls “*simulation.R”* script for every location. “*simulation.R”* reads in the imputed climate data files, and calculates derived climate measures (cumulative cues, fitness, etc). It then calls *“rate-setups.R”* to initialize mutation rates and starting trait ranges. *“simulation.R”* then uses the function `year_var_analyze()` to calculate climate metrics, which are saved in “…-stats.Rdata” in the data-years folder. The script then saves the current scripts (into the results folder), parameters (into the results folder), and the random order of years to be used (into the yearinds folder). These are useful for tracking changes to the model as well as making comparisons between model runs. Climate change scenarios are then generated, creating a changed version of the locations climate data, which is saved in the results folder. *“simulation.R”* then uses parallel processing (using between 1 and 7 clusters depending on the computer’s number of cores) to carry out the actual simulation, which is run through the `run_sim()` function. This and most other functions are defined in the “*windows-subs.R”* file. Results of each simulation run (along with intermediate objects and functions) are saved in the corresponding results folder in a file labeled *“dat.Rdata”*. The realized relative cue use is then calculated and saved in *“finalpop-yrtest-acteff- dat.Rdata”* (see Methods for an explanation of this metric). As a reminder, this is calculated for each individual of the final population exposed to each year of the historic climate. The climate change scenario is then evaluated using the `climate_master()` function. This function takes the final population of the simulation and evaluates its emergence and performance on each year of the changed climate, saving the results in “*temp.Rdata*” in the climate folder within the corresponding results folder. Finally, fitness parameters used for this location are saved in “*…_fitparms.Rdata*” in the corresponding results folder.

After each simulations for each location have been run, “*master-script.R”* uses `trait_eff_summarize_small()`to aggregate the realized relative cue use information to the individual (“*acteff-agg-byindiv.Rdata”*) and the simulation level *(“acteff-agg-bysim.Rdata”*) level in the corresponding results folder, and aggregates all simulation information to the simulation level in *“all- finalpop-yrtest-simlevel-….Rdata”* in the appropriate results folder. After all simulations and aggregations have been carried out, “*master-script.R”* calls `dat_sum_merge()` to summarize climate metrics, and then merge these metrics with location data, simulation results, and realized relative cue use in historic and changed climate regimes, which is saved in into a single data frame with simulation-level summaries which is saved in *“climateVsPops-….Rdata”*. This file contains the majority of the information presented in this paper. Below is a description of each column’s contents.

In these descriptions and in the parameter guide that follows, “simulation” refers to one of several identically parameterized instances of the model in a location; these are best imagined as alternate realities. The results we present had 30 simulations for each location for each set of parameters. “Runs” refers to different executions of the code (typically an execution of the code applies the parameters to every location, producing multiple simulations for each of 78 locations). Presumably these runs are carried out with the purpose of comparing consequences of differences in parameters; thus each run is likely to have a different parameter file.

The results file has a row for each simulation. Each simulation in a location had a different random sequence of years (1000 by default) drawn from the same original collection of years of recorded daily climate data with missing values imputed (see Environmental Data in Methods and Supplements). Unless stated otherwise, climate metrics were calculated based on the sequence of years actually experienced, producing slightly different values between simulations.

### Results file column contents

- name: Location ID (see supplements table 1)
- sim: simulation number
- eff.day: simulation day effect, averaged across all individuals and all years using the *acomp* function in the *compositions* package (see realized relative cue use in Methods)
- eff.cutemp: as day, for cumulative temperature
- eff.cuprecip: as day, for cumulative precipitation
- emerge: emergence day of final population averaged across all years and all individuals
- different values between simulationemerge.cc: emergence day of final population under climate change scenario averaged across all years and all individuals
- geofit: raw units of “fitness” obtained by final generation (used to determine relative). Averaged across all individuals in all years
- geofit.cc: as geofit, but using climate change climate data
- lis.tpred: climate metric that represents the predictability of temperature^5^. This was calculated for the original sequence of years (for each location, the years were ordered chronologically and treated as sequential). As such, the same value was used for all simulations with the same location.
- lis.ppred: as lis.tpred, but for precipitation
- lis.mpred: as lis.tpred, but for moisture
- lis.tseason: climate metric that represents the seasonality of temperature^5^
- lis.pseason: as lis.tseason, but for precipitation
- lis.mseason: as listseason, but for moisture
- pau.tvar: climate metric that represents variability in temperature^6^
- pau.mvar: as pau.tvar, but for moisture
- pau.pvar: as pau.tvar, but for precipitation
- pau.tseason: climate metric associated with each simulation. This metric represents seasonality of temperature^6^
- pau.mseason: as pau.tseason, but for moisture
- pau.pseason: as pau.tseason, but for precipitation
- b.day.mn: day trait of final population, averaged across all individuals
- b.cutemp.mn: as b.day.mn, but cumulative temperature
- b.cuprecip.mn: as b.day.mn, but cumulative precipitation
- med.cv: coefficient of variation in the “median day”. For each year of the climate data, we determined the first day the cumulative temperature reached or exceeded half the maximum cumulative temperature of the mean year for that location. Coefficient of variation was calculated from these median days
- imput.temp.mn: fraction of the daily temperature values that originated form imputation
- imput.prec.mn: as imput.temp.mn, but for precipitation
- med.var: as med.cv, but variance instead of coefficieint of variation.
- within.mn: The “within-year unpredictability” was calculated for each year by comparing the daily temperature to the daily temperatures of the mean year. Each year was allowed to advance or retreat by discrete days, and the temperatures could increase or decrease by a constant; using sum of squared errors, we found the daily shift and temperature change constant to make the current year best match the average year. The “within-year unpredictability” was then the mean of the square of the lag 1 difference (“diff”) of the residuals of this fit. This represents day-to-day inconsistencies that weren’t present in the average year. This metric was calculated for each year of the climate, and then averaged across all years.
- within.nodiff.mn: as within.mn, but using the mean of the square of differences (skipping the “diff” step). This represents divergences between the current year and the average year.
- Var.dailyfit.mn: For each year, we calculated the variance in the relative fitness available on each day. This was averaged across all years.
- lr.var: The “left-right shift” for a year was the number of days advancement needed to best fit the current year’s daily temperatures to the average year of that location. For each location, the lr.var was the variance of these measures across all years for that location.
- Updown.var: as lr.var, but using the temperature shift needed to best fit the current year’s daily temperatures to the average year for that location.
- Temp.mn: mean of all yearly mean temps for the simulation (Note: this was calculated with the temperature values that had been shifted so that the minimum temperature in each location was 0)
- Temp.orig.mn: mean of all yearly mean temps for the simulation using the original, unshifted temperatures
- Temp.predtemp.mn: mean correlation of daily temperature on day n to sum of temperatures on days (n+1):(n+duration). As with all other X.predY.mn, this was a metric for how well X acted as a cue to predict the entire span of Y experienced by an individual that chose to emerge on day n.
- Temp.predfit.mn: as temp.predtemp.mn, but how well temperature predicted sum of fitness
- Temp.predmoist.mn: as temp.predtemp.mn, but how well temperature predicted sum of moisture
- Moist.mn: mean moisture across all years
- Moist.predmoist.mn: as temp.predtemp.mn, but how well moisture predicted sum of moisture
- Moist.predfit.mn: as temp.predtemp.mn, but how well moisture predicted sum of fitness Moist.predtemp.mn: as temp.predtemp.mn, but how well moisture predicted sum of temperature
- Moist.cv.mn: average across all years of the within-year coefficient of variation of daily moisture
- Precip.mn: average across all years of within-year mean precipitation
- Precip.var.btwn: between-year variance in the yearly mean precipitations
- Precip.var.mn: average across all years of within-year variance in precipitation
- Precip.cv.mn: as precip.var.mn, but using coefficient of variation
- Temp.varbtwn: between-year variance in the yearly mean temperatures
- Temp.cvbtwn: between-year coefficient of variation in the yearly mean temperatures
- Precip.cvbtwn: between-year coefficient of variation in the yearly mean precipitation
- Moist.varbtwn: between-year variance in the yearly mean moisture
- Moist.cvbtwn: between-year coefficient of variation in the yearly mean moisture
- Temp.lag1temp.mn: mean lag 1 autocorrelation of daily temperature. As with all X.lag1Y.mn metrics, this was a simple metric for how well X acted as cue to predict Y
- Temp.lag1moist.mn: mean correlation between temperature of day n and moisture of day n+1
- Precip.lag1precip.mn: mean lag 1 autocorrelation of precipitation
- Moist.lag1temp.mn: mean correlation between moisture on day n and temperature on day n+1
- Moist.lag1moist.mn: mean lag1 autocorrelation of moisture
- Moist.mn.var: same as moist.varbtwn
- Temp.cv.mn: mean across years of the within-year coefficient of variation of daily temperatures
- Temp.varwin.mn: as temp.cv.mean, but variance instead of coefficient of variation
- Moist.varwin.mn: as temp.varwin.mn, but moisture instead of temperature
- Lat: latitude of location
- Lon: longitude of location
- Elev: elevation of location
- Coast.dist: distance from location to nearest coast
- Count.good: number of years of climate data available.

### Parameter file

The specifics of any simulation run are controlled by the parameters defined in the parameter file. Below are descriptions of each parameter

- runs.types: the name of each location file to use, in vector of characters
- traits: used for sensitivity analysis. Changing from “day”, “cutemp”, and “cuprecip” may cause the current code to break
- num.sims (default: 30): the number of simulations to run for each location. Each simulation has identical parameterizations, but the randomly generated starting genotypes, randomly generated sequence of years, and randomly generated mutations of each generation will differ.
- N (default: 500): number of individuals per simulation
- num.years (default 1000): number of years to simulate.
- duration (default: 10): number of days organism gathers fitness
- lag (default: 1): number of days after organism decides to emerge before it begins gathering fitness
- mut.dist (default: .005): fraction of trait range to use as standard deviation of mutation rate
- fattail (default: FALSE): indicator to determine whether to use a normal distribution or Cauchy distribution to generate mutation values. Early testing showed that the two distributions produced indistinguishable dynamics in the model, and subsequent simulations used the normal distribution. Setting this to TRUE may cause the current code to break.
- base.temp (default: 0): threshold below which temperature isn’t added to cumulative temperature
- decay (default: .8): decay parameter alpha for use when calculating moisture (see Methods)
- fit.shape (default: “skewgauss”): defines which fitness shape script to use. The “*skewgauss.R*” is the only fitness shape script designed to work with the current code.
- Other.name (default: “moist”): determines what daily environmental measure combines with temperature to define fitness.
- shape.temp (default: 15): initial guess at shape parameter for the temperature fitness curve. The final shape parameter is determined by minimizing sum of squared errors.
- shape.other (default: 15): as shape.temp, but for the other fitness-determining environmental measure (by default, moisture)
- best.temp.quant (default: .9): Used to fit fitness curves. At this quantile of temperatures for each location, the skew normal has its maximum value (see Methods)
- best.other.quant (default: 0.9): as best.temp.quant, but for the other measure (by default, moisture)
- min.quant (default: 0.1): Used to fit fitness curves. At this quantile of temperature and moisture, skew normal function has a value that is (ratio.min) of the peak. (see Methods)
- ratio.min (default: 0.1): determines ratio of values at the min.quant value and best.*.quant value for each skew normal curve.
- plot.extra (default: FALSE): If true, plot emergence of a subset of the generations of each simulation
- plot.pheno (default: FALSE): If true, plot phenotype of a subset of the generations of each simulation
- burnin (default: 100): Used for emergence and phenotype plots, defines how many of the initial generations to skip in the plots
- year.label (default: “A”): When the simulation script is generating a random sequence of years for all simulations in a given location, it checks to see if there exists a file storing random sequences for this location with the same num.sims, num.years, year.set (below) and year.label; if it finds such a file, it uses those randomly generated year sequences. That is, no two simulations within a run will have the same sequence of years, but using the same year.label, num.sims, num.years, and year.set values when running two different parameter files will re-use the same num.sim (default: 30) randomly generated sequences of years for each of the separate parameter file runs. This allows for fair comparisons between different model structures or parameterizations (since ABERDEEN.MS run 1 of each parameterization will have the same sequence of years). Changing the year.label between parameter files causes each to have a separate sequence of years.
- year.set (default: “all.years”): as year.label – shows up in a different part of the naming scheme but plays the same role.
- temp.increase (default: 0 for “shift” climate change, 3 for “warming” climate change): how much should each daily temperature be increased in the climate change scenario?
- precip.shift (default: 5 for “shift” climate change, 0 for “warming climate change”): how many days should precipitation be advanced in the climate change scenario?
- Lockstep (default: TRUE): should temperature advance with precipitation in the climate change scenario?
- tempsd.increase (default: 1.2): for use in deprecated climate change scenario
- phensd.increase (default: 1.2): for use in deprecated climate change scenario
- intra.increase (default: 1.2): for use in deprecated climate change scenario
- scen.temp (default: TRUE): with change in climate change functions, this should always be TRUE
- scen.tempsd (default: FALSE): with change in climate change functions, this should always be FALSE
- scen.phensd (default: FALSE): with change in climate change functions, this should always be FALSE
- scen.intra (default: FALSE): with change in climate change functions, this should always be FALSE
- save.small (default: TRUE): If FALSE, saves information on every individual of every generation in each simulation. If TRUE, only saves information on every individual of the first and final generations.
- photo.flag (default: FALSE): if TRUE, use cumulative photoperiod instead of day of year for the “day” cue.
- do.lisovski (default: FALSE): if TRUE, calculates metrics based on Lisovski et al.^5^, and saves them in separate data files in data-years-lisovski. As this is a lengthy calculation, it’s best to re-use existing calculations of this (and the data files of these calculations can be found in **X**).
- stats.lisovski: if TRUE, uses the metrics associated with do.lisovski. This should be TRUE unless you (a) are missing the lisovski metric data files, (b) don’t want to wait the additional hours for the scripts to calculate them, and (c) don’t want to use those metrics.

